# Virtual Reality and Tablet Cognitive Training Improve Attention and Academic Skills Without Dose Effects

**DOI:** 10.1101/2025.09.16.676629

**Authors:** Anastasia Giannakopoulou, Ariel M. Gordon, Courtney E. Gallen, Michael Seaman, Dominick Fedele, Joaquin A. Anguera

**Affiliations:** Neuroscape Center, Department of Neurology, University of California, San Francisco, United States of America; Department of Psychiatry, University of California, San Francisco, United States of America; Mastermind Cognitive Training, United States of America; School of Sport, Psychology and Social Science, University of Bedfordshire, Luton, United Kingdom; MRC Cognition and Brain Sciences Unit, University of Cambridge, Cambridge, United Kingdom

**Keywords:** attention, video game, cognitive control, executive function, virtual reality, digital

## Abstract

We evaluated the cognitive and academic effects of a closed-loop video game delivered in virtual reality (VR) and tablet formats, at two different dosages, in a school-based setting. A total of 158 children aged 8-9 with a range of attention abilities completed 30 training sessions over 10 weeks. Compared to an expectancy-matched control group, both VR and tablet training led to significant improvements in teacher-rated inattention, performance-based attention tasks, and eye-tracking measures. While both VR and tablet versions of the intervention showed benefits on specific attention-related and academic outcomes, the VR version showed select advantages in both regards. Notably, intervention dose did not significantly moderate outcomes, suggesting that efficacy may depend more on reaching a threshold of engagement than on total duration. These findings demonstrate the utility and benefits of using each type of technology to enhance measures of cognitive and academic abilities as part of a regular school curriculum.

## Introduction

Cognitive control abilities (e.g. attention, working memory, goal management) develop rapidly across childhood, directing our ability to learn and accomplish selected behavioral goals^1,2^. These abilities are especially critical for academic achievement, self-regulated learning, and broader developmental outcomes^3,4^. Deficiencies in these processes, ranging from sub-clinical struggles to attention deficit hyperactivity disorder (ADHD), are prevalent in school-aged populations and contributing to widening disparities in educational attainment^5,6^.

Traditional interventions, such as behavioral classroom strategies, one-on-one tutoring or pharmacological treatments, can yield benefits but are often constrained by variability across teachers, cost, scalability, and limited ecological validity in school settings^7^. The implementation of cognitive neuroscience principles into digital technologies that incorporate real-time task adaptivity and gamification approaches have driven the design of targeted cognitive training programs aimed at enhancing such abilities. Digital cognitive training interventions have been successfully used to enhance deficient cognitive control abilities across a variety of populations^8–21^, including children with inattention concerns^9,22–24^. Such approaches have led to gains in primary schoolers’ attention^25,26^ and inhibitory control processes^27–29^, as well as academic outcomes including reading and mathematics^30–33^. These interventions are often administered with tablet computers due to their portability and ease in facilitating flexible training in naturalistic settings.

However, training on tablets can be challenging in a distractor-rich environment such as typical classroom during a school day. One alternative that has only begun to emerge in recent years is the use of immersive Virtual Reality (VR) to address such limitations by enabling controlled yet richly interactive simulations that limit external influences. Meta-analyses have reported large effect sizes for VR-based interventions in improving attention, memory, and global cognition in children with challenged attention capabilities —typically surpassing outcomes seen in traditional modalities^34^. VR training protocols delivered in virtual classrooms have been shown to boost attention and working memory significantly beyond active controls, and show long-term retention that, in some cases, rivals pharmacological interventions ^35–37^. Similarly, recent intervention studies with children diagnosed with ADHD found VR training led to substantial improvements in continuous performance and working memory compared to traditional interventions^37,38^. It has been demonstrated that such gains in these core cognitive skills are associated with improved academic outcomes^39,40^, although effect sizes are often variable and sensitive to training context and dosage^41^.

Dosage considerations—referring to duration, intensity, and frequency of intervention exposure—remain central to the efficacy of cognitive training programs, particularly in developmental populations. Despite the proliferation of digital and gamified interventions targeting attention and other cognitive abilities, there is limited understanding on how much training is required to elicit robust or lasting effects^42,43^. Furthermore, studies have highlighted a nonlinear relationship between training dose and cognitive outcomes, with diminishing returns beyond moderate levels of engagement^44^. In VR-based interventions, the immersive nature of the modality have been thought to facilitate deeper cognitive engagement per session^45^, potentially reducing the number of sessions required compared to traditional screen-based approaches^46,47^. Yet, high-intensity interventions risk inducing fatigue or disengagement, particularly in younger children. As such, optimizing dosage involves balancing cognitive challenge, motivational design, and developmental appropriateness.

These findings support the assertion that serious games and immersive digital media can be potent levers for the enhancement of attention and overall cognitive performance in children. Yet questions remain about their real-world efficacy, their optimal delivery format, and dosage requirements. In this study, we looked to address each of these open questions through an in-school delivered, randomized controlled trial using a digital, neuroscience-informed intervention designed to enhance cognitive performance versus an expectancy-matched placebo control. The intervention included adaptive exercises targeting cognitive functions such as attention, inhibition, visual tracking, and working memory. The study was deployed across diverse school settings using two formats, immersive VR and tablet, with its design aimed to evaluate three key hypotheses:

Hypothesis (1) participants who received the intervention would show greater attention-based improvements than those in an active expectancy-matched control group; Hypothesis (2) participants who trained using the VR format would demonstrate greater gains than those using the tablet format; Hypothesis (3) participants assigned a higher training dosage (e.g. 25 minutes for each training session as opposed to a ‘low dose’ 12.5 minute session) would show greater gains across all training outcomes.

## RESULTS

### Overview of Study Design and Demographics

A total of 168 children (between 8-9 years of age, see **Table 1**) attending 3 different elementary schools near Luton in the United Kingdom were presented with the opportunity to participate in a research study designed to address Hypotheses 1-3. All participants were given an overview of the study requirements as well as consenting information to be signed by their parents, as well as assents signed by the participants. Enrollment was done at the class level (see Methods and CONSORT diagram, **Figure 1**, for more details), with all children given the opportunity to play a custom digital intervention created by Mastermind

**Figure 1.**
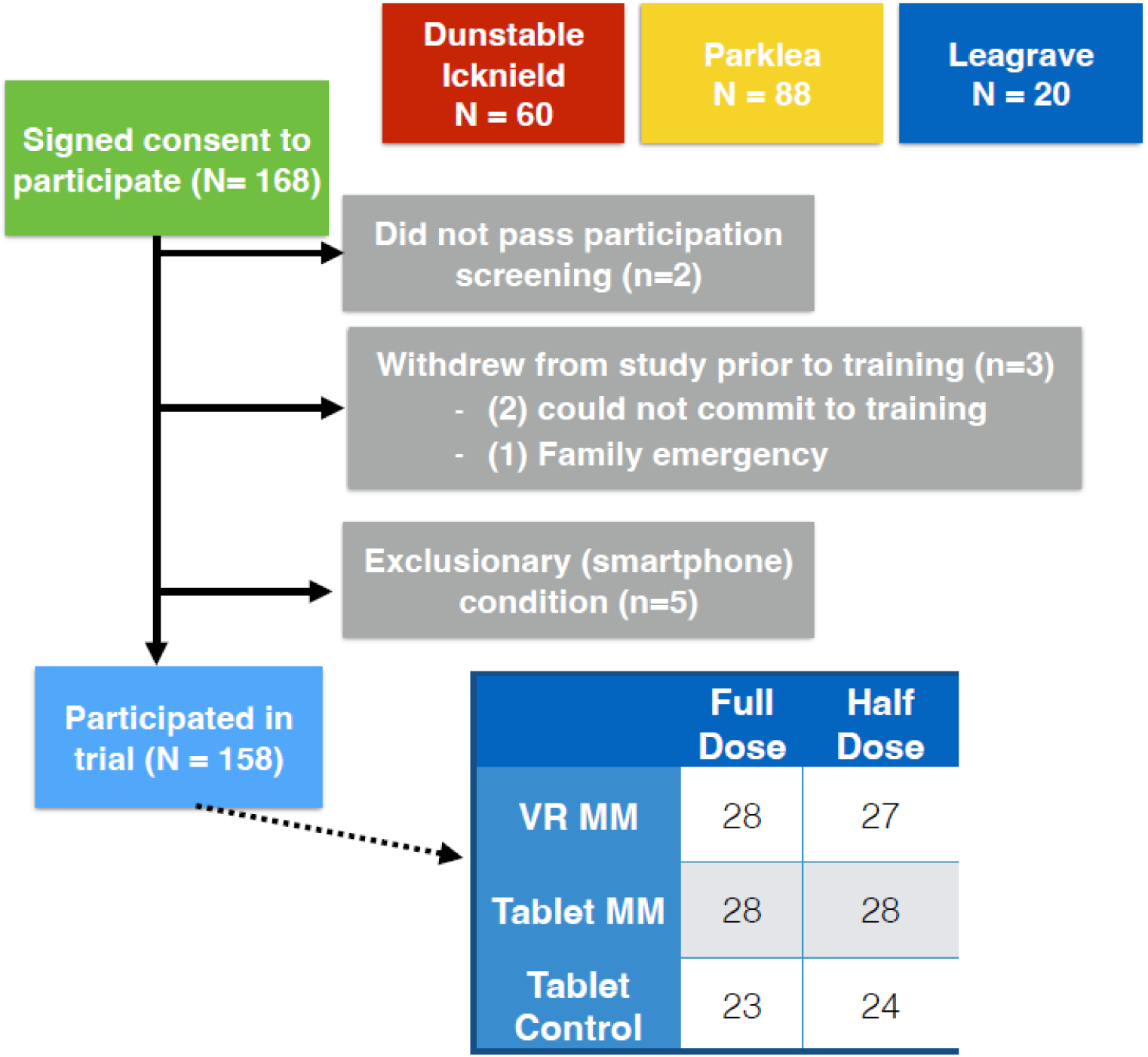
CONSORT Diagram

**Table 1.** Baseline demographic characteristics of the participant sample. The continuous variables are summarized as the mean (standard deviation).

| Demographics | VR<br>(full dose) | VR<br>(half dose) | Tablet<br>(full dose) | Tablet<br>(half dose) | Control<br>(full dose) | Control<br>(half dose) | Non-Participant | Total |
| --- | --- | --- | --- | --- | --- | --- | --- | --- |
| Number of<br>participants | N= 28 | N= 27 | N= 28 | N= 28 | N= 23 | N= 24 | N=10 | **N= 168** |
| Age | 8.64<br>(4.21) | 8.74<br>(3.39) | 8.72<br>(4.92) | 8.71<br>(4.45) | 8.71<br>(3.64) | 8.64<br>(4.69) | 8.71<br>(2.92) | **8.71**<br>**(4.15)** |
| Sex | 14m, 14f | 11m, 16f | 17m, 11f | 13m, 15f | 12m, 11f | 14m, 10f | 6m, 4f | **87m, 81f** |

Cognitive Training™ or a battery of freely available math and coding applications (Kahoot!, Sushi Monster, Dance Party, ScratchJr) that acted as the control arm of this study, with participants in the intervention and control group given either a full dose of daily training (∼25min, ‘Full’) or a half dose (∼12min; ‘Half’). The Mastermind intervention was presented in virtual reality (via Oculus Quest 2, Meta™) or tablet (iPad, Apple™; see **Figure 2a-c**), while the control was presented only in tablet form (see **Figure 2d**). Students in each class were randomized to one of the three possible interventions using a block design with a block size of six to maintain balanced group sizes. The dosage prescribed was the same for each class for practical purposes (e.g. to ensure that all student trainings ended at the same time in said class for continuity with their school day). Participants trained in groups of five per session in separate rooms, and under the supervision of at least one research assistant per group to monitor participation, deliver task instructions, and provide technical support for the equipment in use. Students completed 30 training sessions over 10 weeks during the school day at a consistent class time each week; make-up days were provided each week to allow those students who missed a training day to catch-up on their training sessions. Teachers at school were blinded as to which intervention and dosage were being given to their students, with research assistants present for each assessment and training session. Participants who reached the training portion of the study completed 100% of all assigned training sessions.

**Figure 2.**
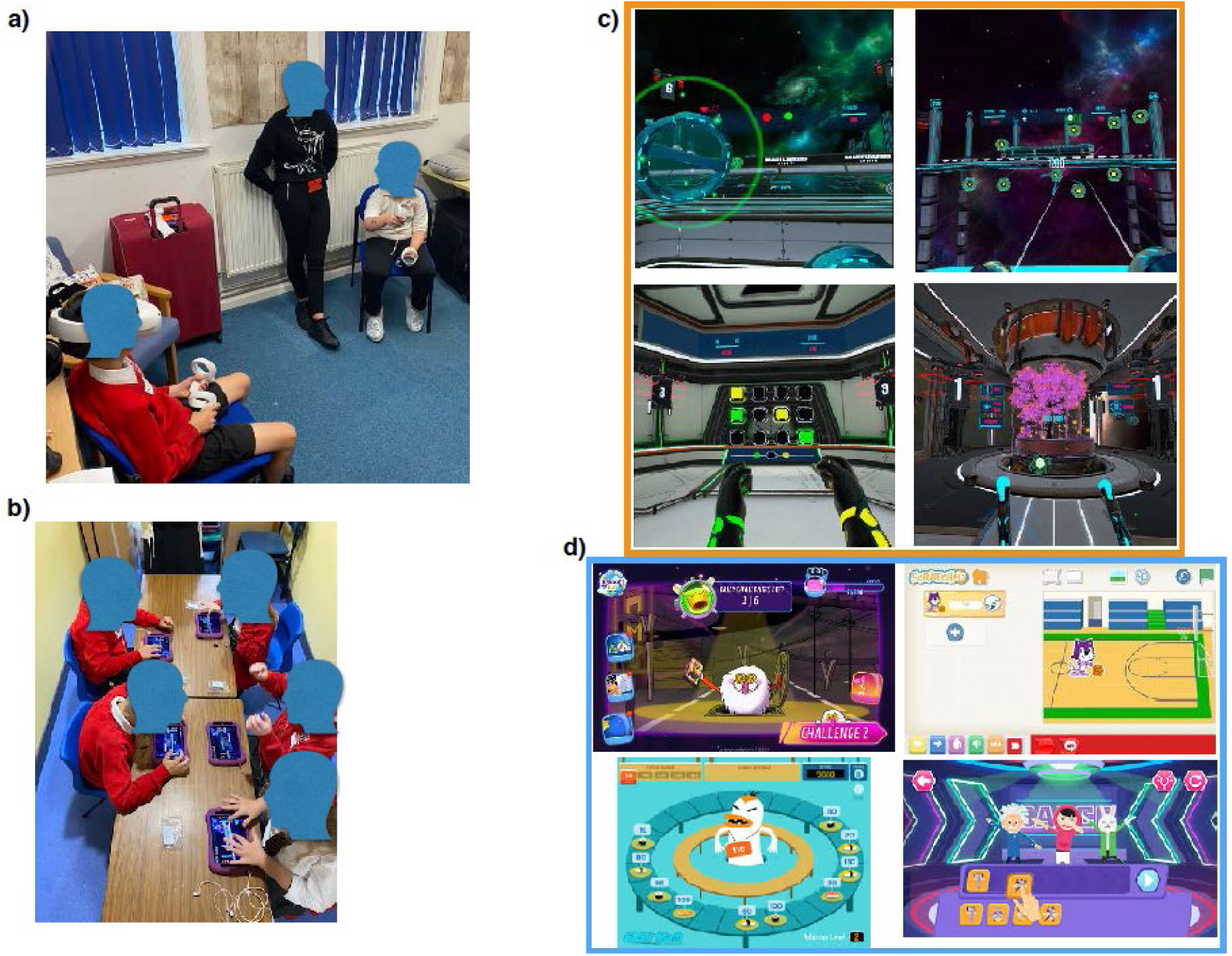
Images of training setting and platforms. Additional details and images of the training program can be found at mastermindtraining.com, with videos of the VR tasks shown at (https://youtu.be/pJ0IPutvPMU).

*Expectancy:* After the 1^st^ day of training, students were asked to complete an expectancy survey to evaluate their impressions of how much improvement they expected on the objective outcome measures of attention (Right Eye, mobile continuous performance task (mCPT); see **Methods** for more detailed description), a custom test of reading and math fluency (SEA^20^), and how enjoyable they believed their assigned intervention would be. Four separate analysis of variance (ANOVA) tests with the factor of group (6 groups; Full VR MM, Half VR MM, Full Tablet MM, Half Tablet MM, Full Control, Half Control) were used to test for perceived differences on each outcome measure (Right Eye, mCPT, SEA math and SEA reading, ‘fun’). There were no significant group differences on the perception of improvement across groups on the Right Eye measure (F_(5,149)_= 1.69, p= 0.14), the mCPT measure (F_(5,149)_= 1.59, p= 0.16), SEA math fluency (F_(5,149)_= 1.58, p= 0.17), SEA reading fluency (F_(5,149)_= 1.29, p= 0.27). However, there was a slight group difference with respect to the perceived enjoyment regarding assigned intervention (F_(5,149)_= 2.31, p= 0.47). When collapsing across dose and technology (e.g. all Mastermind participants versus all control participants), follow-up tests revealed that there was no difference in perceived ‘fun’ (t_152_ = .48, p= 0.63) between these two groups. Further testing of this effect revealed that while neither the VR MM or Tablet MM interventions were perceived as being more ‘fun’ than the control (t_(60.2)_ ≦1.57, p≧0.12 in each case), the VR MM was perceived as being more ‘fun’ than the MM Tablet intervention (t_(66.5)_ =2.76, p= 0.007).

### Outcome measures

*Right Eye:* Here we assessed whether the MM intervention led to enhanced post-training overall eye-tracking performance beyond that of the control (Hypothesis 1). Using an analysis of covariance analytical approach (ANCOVA; see **Statistical Analysis** for more details), we observed a greater overall Right Eye score following the training period by the MM intervention group than that of the control group (F_(2,150)_= 11.2 p= 0.001, **Figure 3a)**. Given that the overall Right Eye performance metric is constructed of a combination of fixation, saccades, and pursuit eye movement measures, we subjected each of these individual post-performance metrics to the same ANCOVA analysis to interrogate which measure(s) were driving the overall post-training effect (as described in our trial registration). This testing revealed that the MM intervention showed enhanced post-training performance beyond the control group on measures of fixation (F_(2,150)_= 4.36 p= 0.04), and saccade (F_(2,150)_= 8.70 p= 0.004), but not pursuit (F_(2,150)_= 2.36 p= 0.13; see **Supplementary** Figure 1a-c).

**Figure 3.**
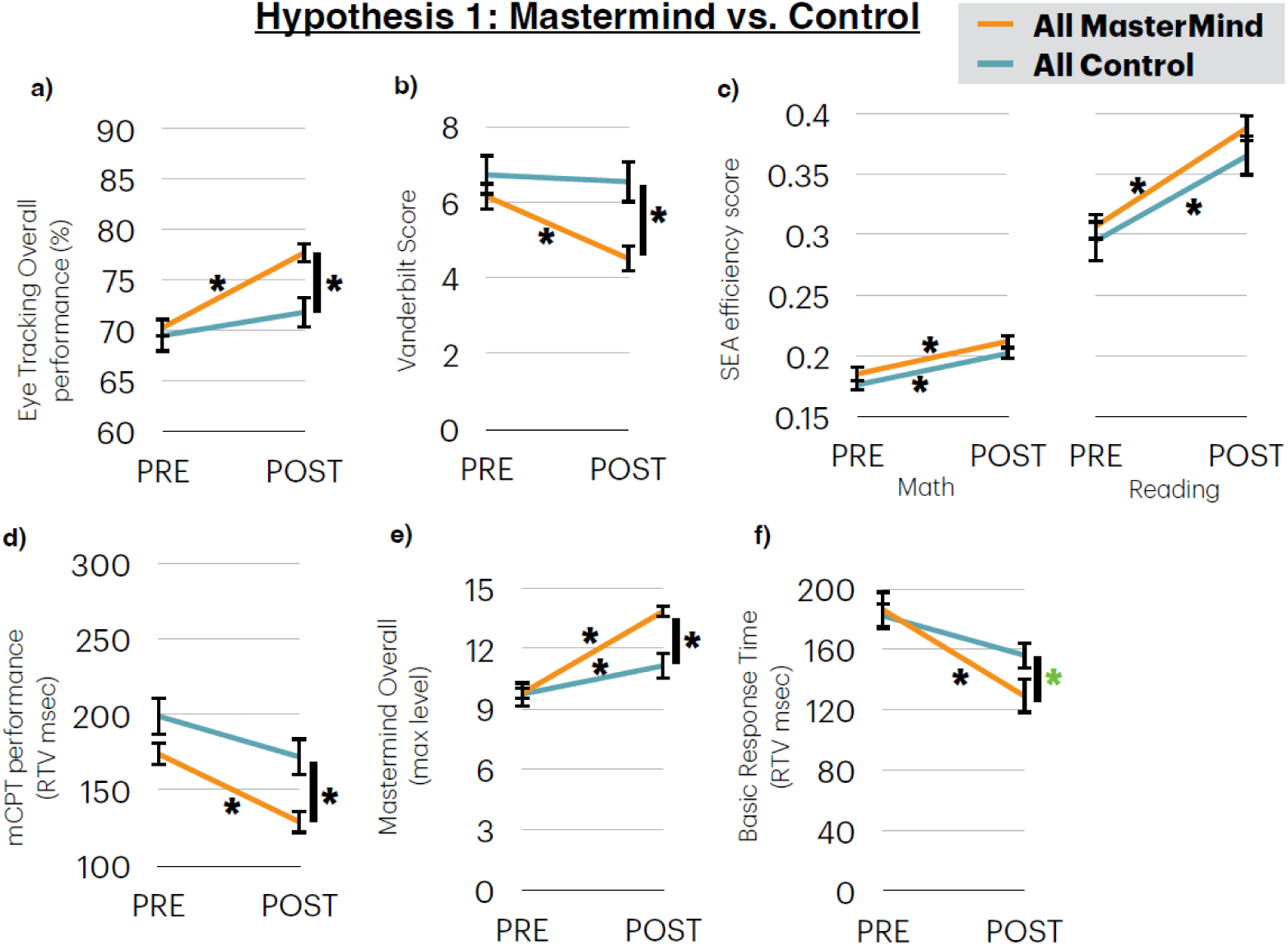
Panels of each group comparison for Hypothesis 1 for each measure. * = p< .05, **|* =** significant ANCOVA. Green * p < .07 (trend).

When we assessed Hypothesis 2, we observed no difference between the VR and Tablet MM groups on their overall eye-tracking score (F_(2,104)_= 2.72 p= 0.10, **Figure 4a**). Interrogating this further by examining fixation, saccades, and pursuit eye movements measures as in Hypothesis 1, this analysis revealed that the VR MM intervention showed enhanced post-training performance beyond the Tablet MM group only on the measures of pursuit (F_(2,104)_= 7.67 p= 0.007), but not for fixation (F_(2,104)_= 0.14 p= 0.91) or saccades (F_(2,104)_= 0.30 p= 0.58; see **Supplementary** Figure 2a-c).

**Figure 4.**
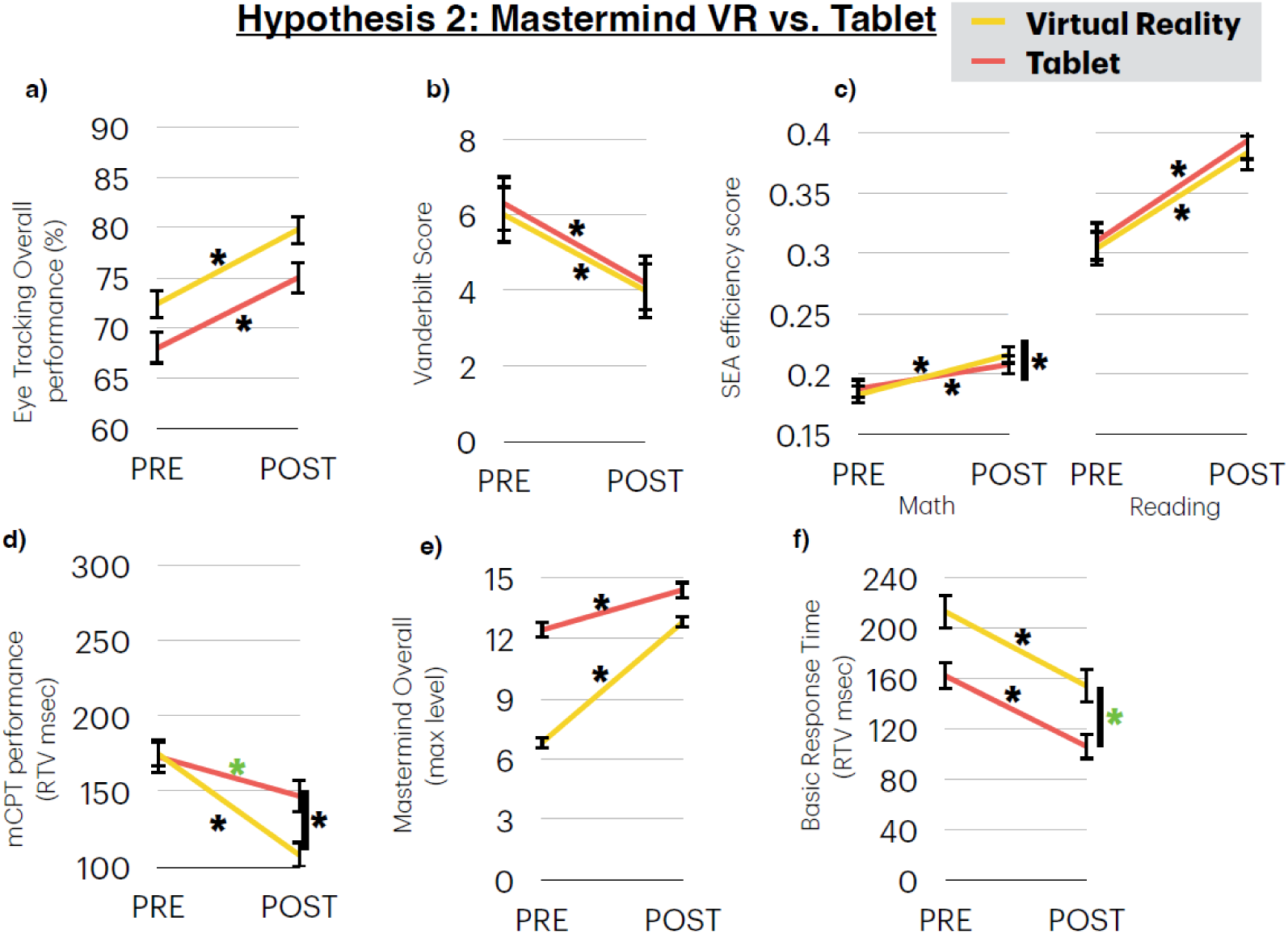
Panels of each group comparison for Hypothesis 2 for each measure. * = p< .05, **|* =** significant ANCOVA. Green * p < .07 (trend).

Finally, we used the same analytical approach to test whether the dosage of training across the VR MM and Tablet MM groups differentially impacted the post-training eye-tracking scores (Hypothesis 3). Here we observed no significant dose by group interaction on their overall post-training eye-tracking scores (F_(4,102)_= 1.47 p= 0.23, **Figure 5a**). However, following the same interrogation of the individual tests as in the Hypothesis 1 and 2, we did observe a significant dose by group interaction for fixation (F_(4,102)_= 8.24 p= 0.005), but not for pursuits (F_(4,102)_= 0.46 p= 0.83) or saccades (F_(4,102)_= 0.10 p= 0.75). Follow-up tests using the same ANCOVA approach interrogating each potential group pairing revealed that the VR full group showed a significant difference from the VR half group (F_(2, 49)_= 7.60 p= 0.008) and a trend towards a VR full advantage over iPad full (F_(2, 51)_= 3.51 p= 0.067), with all other comparisons not showing any group differences (F≤ .98 p≥ 0.32; see **Supplementary** Figure 3a-c). Group means and standard errors are presented for both pre-training and post-training for all outcome measures for each hypothesis in **Supplementary Tables 1-6**.

**Figure 5.**
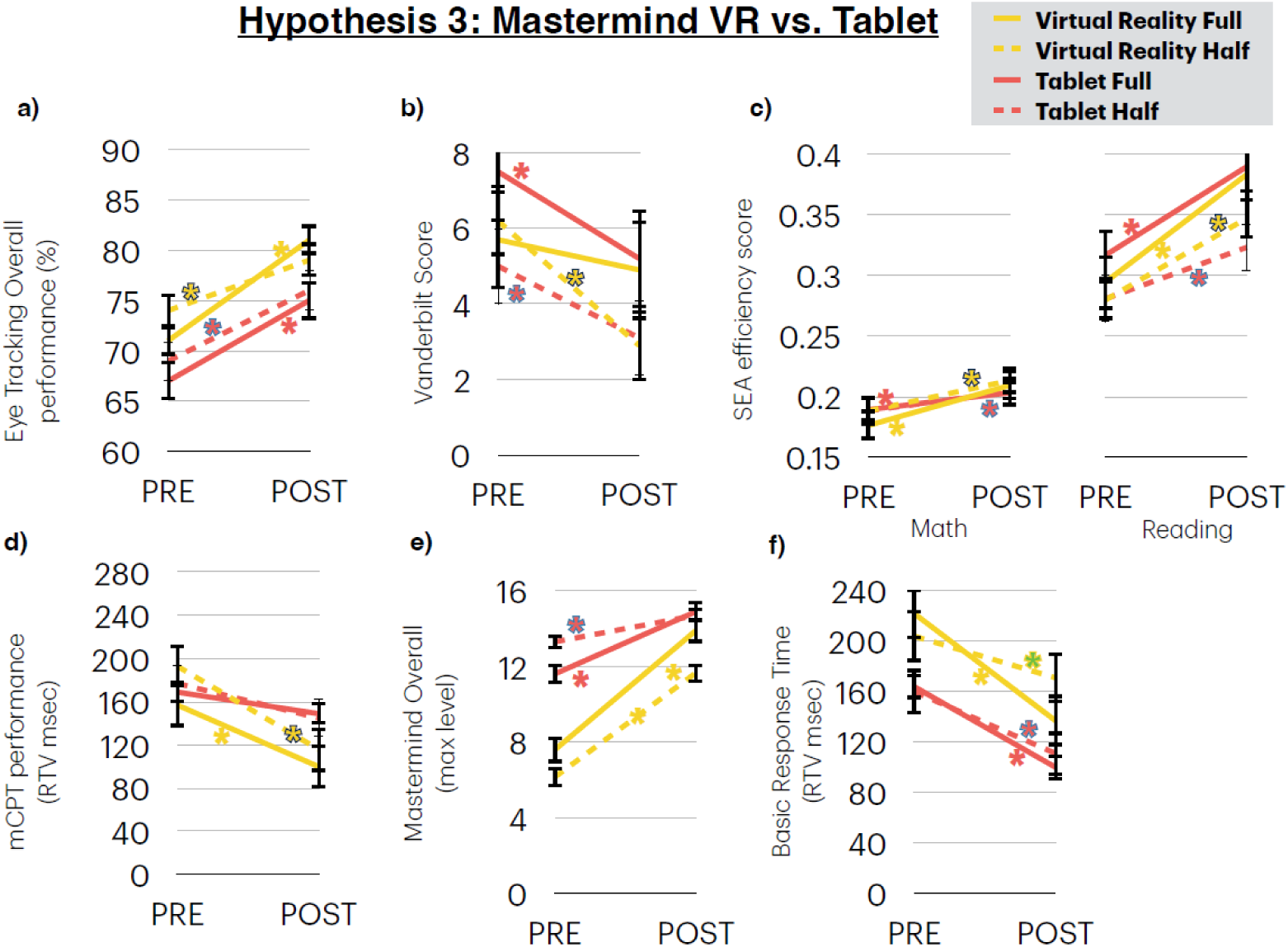
Panels of each group comparison for Hypothesis 3 for each measure. * = p< .05, **|* =** significant ANCOVA. Green * p < .07 (trend).

*Vanderbilt:* To test whether the MM intervention led to changes in teacher perceptions of inattention beyond that of the control (Hypothesis 1; with teachers blinded to group assignments), we examined the post-training teacher reported scores on the Vanderbilt measure based on a cumulative total from questions 1-9 on the questionnaire^9^. Using the aforementioned ANCOVA test, we observed a lower inattentive score following the training period by the MM intervention group than that of the control group (F_(2,152)_= 5.87 p= 0.017, **Figure 3b**). To test whether this effect was differentially impacted by the type of training platform utilized by the intervention group (Hypothesis 2), we used the same approach comparing the VR MM and Tablet MM groups (collapsing across dose) and observed no significant difference (F_(2,105)_= 0.12 p= 0.91, **Figure 4b**). Finally, we used the same analytical approach to test whether the dosage of training across the VR MM and Tablet MM groups differentially impacted the teachers post-training inattentive scores (Hypothesis 3). As with Hypothesis 2, we observed no significant dose by group interaction on their post-training teacher reported inattentive scores (F_(4,103)_= 1.25 p= 0.26, **Figure 5b**).

*SEA*: For Hypothesis 1, we assessed whether the MM intervention led to enhanced reading and math fluency that of the control using an ANCOVA approach. We observed no between group difference following the training period for both reading fluency, (F(_2,142)_= .67 p= 0.41, **Figure 3c**) and math fluency (F(_2,144)_= .35 p= 0.56; **Figure 3c**), as both groups showed a significant improvement from baseline in each case. To assess the impact of our active control group, we compared each group’s performance to that of historical control group of similar aged students from another UK school who completed the same SEA assessment following ‘instruction-as-usual’^20^. Using the same ANCOVA approach, we observed that both present groups showed a significant improvement from the students in the Zanto et al. study (see **Table 2**), suggesting that the present control group may have underwent more rigorous training than a typical instruction-as-usual group. We were also presented the opportunity to compare our SEA reading fluency assessment to UK national standard teacher-administered assessments for each student, and found that our SEA measure aligned with these national standards. We present these findings in the **Supplementary materials** as they were not part of our trial registration.

**Table 2.**
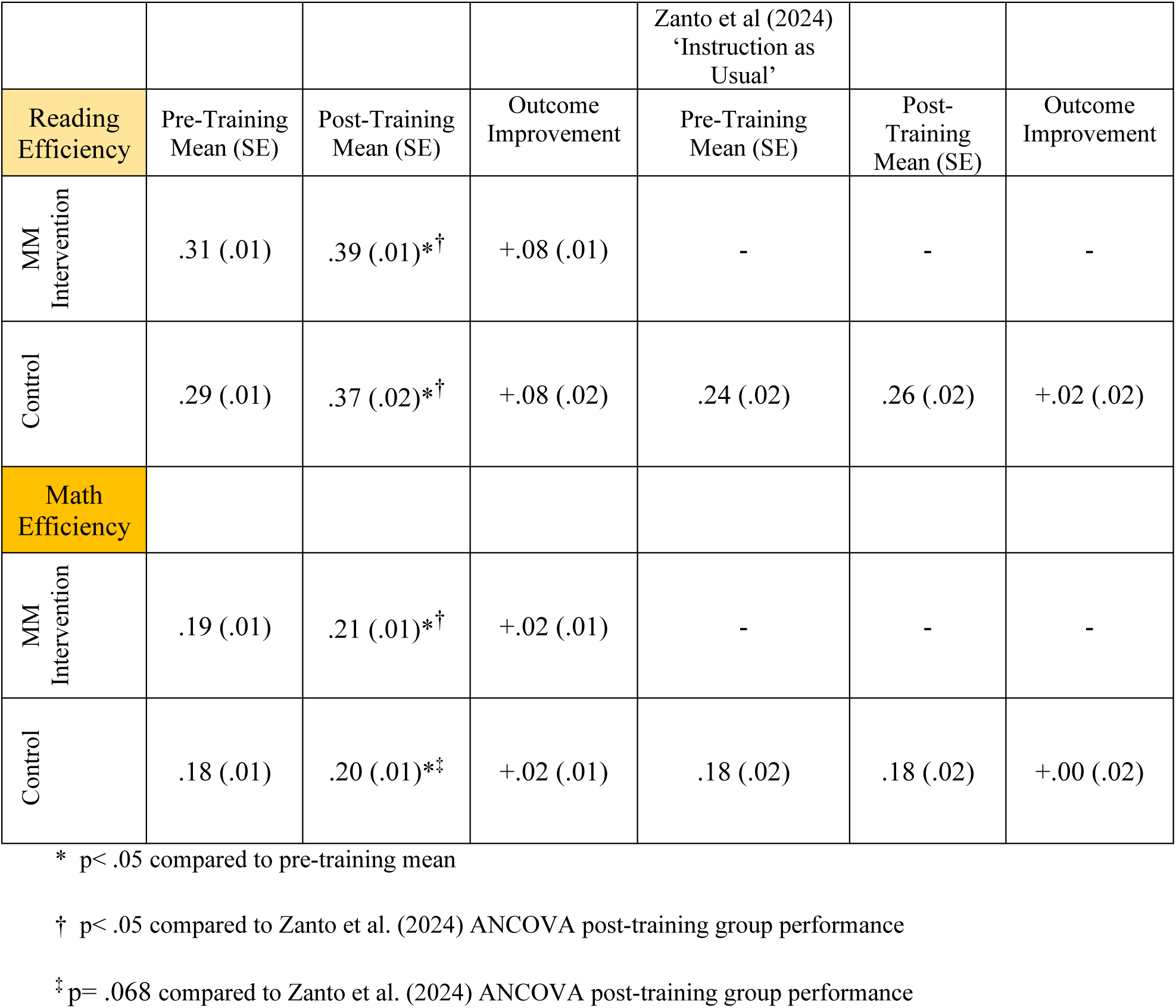
Comparison of SEA Reading and Math Fluency Efficiency to Historical Control.

When we assessed Hypothesis 2, we observed no between group difference in reading fluency (F_(2,99)_= 0.061 p= 0.81, **Figure 4c**), as once again both groups showed significant improvement from baseline but no platform advantage. However, with respect to math fluency we did see an advantage for the VR group beyond the improvement seen for the tablet group (F_(2,100)_= 4.12 p= 0.045, **Figure 4c**). Finally, for Hypothesis 3, the ANCOVA analysis revealed a dose by group interaction for math fluency (F_(4,101)_= 5.10 p= 0.026, **Figure 5c)** unlike reading fluency (F_(4,98)_= .37 p= 0.54, **Figure 5c)**. Interrogation of the significant interaction revealed that only the VR full group showed an advantage beyond the tablet full group (F_(2,49)_= 8.62 p= 0.005), with all other comparisons not showing any group differences (F≤ 2.77 p≥ 0.10).

*mCPT:* To test whether the MM intervention led to improved performance on this measure of attention beyond that of the control (Hypothesis 1), we examined response time variability on the frequent target condition of the mCPT. Using the same ANCOVA approach, we observed improved response time variability following training by the MM intervention group than that of the control group (F_(2,150)_= 8.32 p= 0.004, **Figure 3d**). With respect to whether this effect was differentially impacted by the type of training platform utilized by the intervention group (Hypothesis 2), we observed a significant difference suggesting that the VR group showed a greater improvement than the tablet group (F_(2,104)_= 10.36 p= 0.002, **Figure 4d**). Finally, the test of the dosage (Hypothesis 3) revealed no significant dose by group interaction (F_(4,102)_= .37 p= 0.54, **Figure 5d**).

*Mastermind Assessment:* Here we examined performance on the assessment version of the three different training modules—working memory (Moon Shadow), multiple object tracking (Star Catcher), and task switching (Sun Blazer), using the maximum level achieved in each case, with the average across all three tests contributing to the calculation of the overall MM assessment score. For Hypothesis 1, we observed a greater overall score following the training period by the intervention group than that of the control group (F_(2,152)=_ 46.76 p= 0.0001, **Figure 3e**). Similar to the Right Eye analysis, we subjected each individual module to the same ANCOVA analysis to interrogate which measure(s) were driving the overall effect. This testing revealed that the intervention showed enhanced post-training performance beyond the control group on each measure: working memory (F_(2,150)_= 29.84 p= 0.0001), multiple object tracking (F_(2,150)_= 27.83 p= 0.0001), and task switching (F_(2,150)_= 14.48 p= 0.0001; see **Supplementary** Figure 1d-f). When we assessed Hypothesis 2, we observed no difference on the overall score between the VR and tablet group (F_(2,105)_= 0.23 p= 0.88, **Figure 4e**), nor on any of the individual measures: working memory (F_(2,105)_= 0.010 p= 0.92), multiple object tracking (F_(2,105)_= 1.31 p= 0.25), and task switching (F_(2,103)_= .90 p= 0.34; see **Supplementary** Figure 2d-f). Finally, we did not observe a dose by group interaction on the overall score (Hypothesis 3; F_(4,103)_= 1.65 p= 0.20, **Figure 5e**) or for any individual module: multiple object tracking (F_(4,101)_= 0.42 p= 0.52), working memory (F_(4,103)_= 1.46 p= 0.23) or task switching (F_(4,103)_= 2.72 p= 0.10; see **Supplementary** Figure 3d-f).

*Basic Response Time*: Finally, we administered a measure of basic response time (BRT) to evaluate for changes in variability related to simple motoric quickness or processing speed. Here we observed a strong trend toward a group difference for Hypothesis 1 suggesting an intervention group advantage (F_(2,150)_= 3.87 p= 0.051; see **Figure 3f**). With respect to whether this effect was differentially impacted by the type of training platform utilized by the intervention group (Hypothesis 2), we observed another trend suggesting that the VR group showed less performance variance than the tablet group (F_(2,103)_= 2.97 p= 0.088, **Figure 4f**). Finally, the test of the dosage of training across the VR MM and Tablet MM groups on RTV (Hypothesis 3) revealed no significant dose by group interaction (F_(4,101)_= .77 p= 0.38, **Figure 5f**).

*Correlations:* Here we looked to probe for relationships involving attention-based variables that showed significant group effects for Hypotheses 1 or 2 with our measures of academic performance, with the overarching goal being to explore what form of cognitive training may translate into academic achievement. We observed a correlation between improvements on the mCPT task and reading efficiency gains across all participants (r_(143)_= .22 p= .008; see **Supplementary** Figure 4a). This finding suggests that those individuals that showed the greatest improvement on an objective task of attention were those who improved their reading fluency the most. Given that the intervention group consisted of both VR and tablet participants, when this analysis was performed on each of these groups it was revealed that this relationship was present for the VR group (r_(49)_= .34 p= .018; see **Supplementary** Figure 4b), unlike the tablet group (r_(52)_= .11 p= .44; see **Supplementary** Figure 4c), suggesting a platform specific effect. A similar pattern of results involving the VR group was also seen when using the Mastermind overall score with reading fluency (r_(50)_= .31 p= .029), providing additional support for the proposed relationship involving enhancements in cognitive and reading abilities. Finally, we observed a correlation between the Mastermind overall score with math fluency (r_(103)_= .22 p= .029), supporting the assertion that interventions enhancing cognitive abilities can positively impact skills including numerical fluency.

## Discussion

The present study provides initial evidence regarding the potential benefit that a novel digital intervention platform distributed in both VR and tablet format at different dosages can have on cognitive control abilities, with this trial occurring as part of a regular school day for children 8-9 years of age across 3 schools. These findings demonstrate that this intervention enhanced objective and subjective measures of cognitive control (as well as multiple measures of eye tracking) in school-age children beyond that of an expectancy-matched control. When specifically interrogating these effects across different training technologies, there were few advantages conferred by training in VR beyond tablet. Similarly, when examining the effect of dosage, there were very few outcomes that showed a significant difference between the full and half dosage, providing initial support for considering a shorter, more practical training time in certain scenarios. Here we discuss the implications of each tested hypothesis and how these outcomes align with prior research highlighting the promise of serious games and digital cognitive training in developing cognitive control functions^48^.

For Hypothesis 1, participants using the MM intervention on VR and tablet platforms outperformed an active, control group on multiple measures of attention. This was evidenced by a direct measure of sustained attention (mCPT) and a real world blinded teacher report measure of inattention (Vanderbilt), in agreement with previous digital therapeutics trials that showed comparable enhancements on these measures^10,22,23,49^. Participants that trained with the intervention also demonstrated significantly better eye-tracking performance than controls. These findings are aligned with the present attention results, as oculomotor behaviour has been described as a marker of attentional control^50–54^. Furthermore, multiple studies involving computerized eye-tracking training have also shown to significantly improve attention deficits in children with attentional difficulties^55–57^. The present findings, in conjunction with such studies, provide evidence for the ability to engender neuroplasticity in oculomotor systems following targeted cognitive intervention.

While cognitive control measures were consistently enhanced beyond the control group, our assessment of math and reading fluency did not show a similar between-group differential benefit (that is, both groups improved equivalently). These findings appear inconsistent with previous work where an improvement in reading fluency efficiency was observed using only the Coherence intervention (which is a part of the present MM battery, see Methods) in this same age group versus a group of students that received ‘instruction as usual’^20^. That study suggested that reading improvements could be realized solely through enhanced rhythmic timing ability, as no Coherence-related enhancements on measures of attention, inhibitory control, or working memory were observed^20^. There are key differences between these studies that help explain the lack of group differences on these math and reading measures. Here, the active control group’s improvement in reading fluency was more than 3 times greater than non-significant gain observed by the ‘instruction as usual’ group from the Zanto et al. study. This suggests that the control administered here may have been more effective at improving reading fluency than initially assumed when designing the control arm. The other notable between-study difference involves improvements in cognitive abilities, which were observed only in the present study. There is a deep literature describing the relationship between cognitive functions and academic achievement^58–60^ that supports the idea that enhancing such abilities can benefit reading. Here we observed not only group differences on multiple measures of attention (and working memory via the MM assessment), but we also observed correlations between improvements in reading fluency and both the mCPT and MM assessments. The present findings warrant further investigation to better understand how closed-loop interventions used in conjunction with academic applications could foster even greater benefits. This is especially pertinent given that this cognitive intervention did not have any directed academic focus yet led to comparable academic fluency improvements as applications specifically designed for academics.

This leads to the questions of why would the control battery with math and coding games lead to improved reading fluency, and what is the underlying mechanism driving improved math fluency in the MM platform? Previous work has shown that interventions targeting mathematical skills in children have shown crossover benefits for reading fluency^61,62^, including through the strengthening of cognitive control abilities^63,64^. For example, structured math interventions promoting number sense and arithmetic automaticity simultaneously support phonological processing and decoding efficiency, key factors that are called upon for reading fluency^65,66^. Moreover, interventions emphasizing numerical fluency, such as rapid retrieval of math facts, appear to foster general cognitive fluency that transfers positively to timed reading tasks, suggesting overlapping neural and cognitive resources engaged during math and reading activities^67,68^. Collectively, these findings support the idea that the present control was not only quite rigorous, but also potentially engaged overlapping cognitive processes that improved reading fluency performance. Such conclusions warrant future efforts to not only replicate the improvements in cognitive abilities, but to also better understand how the improvements in cognitive control contribute to these and any other type of math-based enhancements.

Our second hypothesis posited that immersive VR would yield superior gains compared to tablet-delivered cognitive training. This prediction was based on work demonstrating that the VR medium is able to enhance attentional engagement through heightened presence^45,69,70^ as well as reduced distraction, thereby reinforcing attentional control mechanisms that generalize beyond the training environment^71^. Here we observed only a few (but important) advantages when comparing the efficacy of the VR and tablet platforms, with both technologies demonstrating significant improvements from baseline for nearly all measures. The participants training with MM using VR did outperform the tablet group on the mCPT task as well as the MM overall score, with both measures assessing cognitive abilities including attention. Furthermore, only the VR group showed a correlation on each of these measures and reading fluency, suggesting that these two particular VR advantages also impacted reading fluency in a positive manner. The attention-based findings align with meta-analyses involving ADHD populations that specifically examined the use of immersive VR interventions versus non-VR controls on measures of attention (including on CPT measures), with these studies reporting medium to large effect sizes^72–75^. However, an important distinction of the present study from those studies is that it involved children with a range of attention abilities, not just those with ADHD, suggesting that the observed effects here could benefit children across all attentional levels. The VR group also realized greater gains in math fluency—supporting other work demonstrating that immersive platforms can enhance cognitive and educational outcomes beyond touchscreen interfaces^76–79^. Thus, there are certain benefits that came from the use of VR versus the use of tablet technology that may warrant its adoption in certain settings, such as when specifically targeting cognitive control abilities. However, VR is more cumbersome to deploy compared to tablets due to its elaborate hardware, unfamiliar interface, greater physical demands, and limited visibility for administrators. Thus, these initial results warrant replication efforts to verify the utility of incorporating VR technology into a given training program, especially when training on a tablet may suffice.

Finally, our third hypothesis was that the dose of a given intervention would impact the level of efficacy across groups on multiple measures. However, we did not find results to support such a differential effect: the duration of the intervention did not significantly impact the majority of the outcomes. Previous work involving closed-loop digital interventions have shown efficacy with doses that range from as little as 5 hours of training over two weeks^80^, with more typical dosages being between 8-15 hours over the course of 4-8 weeks^8–21^. This finding contrasts with prior meta-analytic evidence suggesting that training duration is a key moderator of cognitive training efficacy, particularly in pediatric populations^81,82^. Previous cognitive training research has highlighted nonlinear dose–response relationships, where gains plateau after an optimal dose, potentially due to homeostatic regulation of plasticity^83,84^. This interpretation is consistent with previous studies suggesting that early performance improvements may plateau after a relatively short period of engagement, especially when task novelty and cognitive demands are front-loaded^85,86^. This suggests that optimal engagement—rather than sheer session duration— in conjunction with specific mechanisms of the intervention including task adaptivity can especially impact training efficacy. Future studies employing individualized dosing and neurophysiological markers of engagement^87^ may better elucidate how closed-loop systems harness experience-dependent plasticity to maximize both domain-specific and generalized cognitive gains.

In conclusion, this randomized controlled trial demonstrated that the present intervention was able to improve attentional control and academic fluency in school-age children. Across both immersive VR and tablet-based formats, the intervention outperformed the active controls on objective attention metrics and teacher-rated inattention, demonstrating the breadth of efficacy and cross-platform versatility. The consistency of effects across modalities suggests that targeted cognitive engagement, not immersion alone, drives behavioral transfer, while immersive VR conferred specific gains that could be associated with academic performance. Finally, the effects observed were largely dose independent aside from specific features where the full dosage of VR had a significant benefit, suggesting that practical considerations of both technologies and available resources should be considered with regards to the outcomes aimed for. These preliminary findings support the use of Mastermind Cognitive Training as a promising, platform-flexible digital therapeutic to address attentional challenges in real-world settings.

One of the primary goals of this study involved the feasibility of incorporating a cognitive training platform in a virtual reality platform in school setting. Here we succeeded in delivering the MM intervention as an in-school activity, with the expectancy findings suggesting that these students enjoyed the training experience, and not just the novelty of using VR. However, there are several practical considerations to consider with respect to replicating these efforts. The required elements here included having a dedicated space that could accommodate a number of students at once, as well as having at least one individual per 5 students to oversee the training protocols. While the space and peripheral equipment needs are not trivial, the possibility of scaling this design appears to be feasible, especially when considering that the costs of these technologies are declining.

However, there are limitations that should be considered with the present findings. First, it is unknown how long the present intervention effects last, as we were unable to collect follow-up data in this trial. Previous work has demonstrated that digital interventions utilizing closed-loop approaches targeting cognitive control abilities have shown that acquired benefits can last anywhere from 6 months to up to 3 years following the initial training period^8,9,12,21,22,24^. Follow-up work that assesses the persistence of these effects on both measures of attention and academic metrics would be especially intriguing, or if booster sessions following the initial training period may be optimal for long-term effects. Second, the study design lacked a virtual reality control group (with each dosage), necessitating a between-platform comparison. This concern is mitigated in part by the expectancy results demonstrating that the VR and control groups had similar perceptions regarding the amount of improvement expected and enjoyment to be had. Third, while the present study utilized research measures assessing cognitive abilities as well as reading and math fluency, these are not the same as established benchmarks used by teachers to assess academic achievement.

While we were able to interrogate certain measures used by teachers, these findings should be considered exploratory given that these analyses were not a part of the trial registration.

Finally, it is unclear to what extent the benefits observed here in 8-9 year olds would be observed in other groups, including older students (e.g. teenagers), older adults, or individuals with ADHD. Given the breadth of benefits that other closed-loop platforms have had in other populations across multiple ages^8–21^, including the present platform with expert tennis players^88^, the present findings support future work examining its impact across the lifespan and clinical conditions.

## Methods

The study design, data collection, and statistical analysis plan of this randomized control trial was registered on the ISRCTN registry with the number ISRCTN75915439 (https://doi.org/10.1186/ISRCTN75915439). The study was approved by the University of Bedfordshire Research Centre for Applied Psychology (RCAP) Research Ethics Committee (REC) (approved protocol reference RCAP072024). The study was conducted in accordance with the Department of Psychology, University of Bedfordshire policies and regulations, following the British Psychological Society Code of Ethics and Conduct^89^ and the Declaration of Helsinki. Written consent was provided by the primary caregiver, with participants providing informed assent prior to participation. They were informed that their participation was strictly voluntary and that they had the right to withdraw their participation at any time during the data collection period. Prior to obtaining consent, participants were confirmed to have the cognitive capacity to provide assent by the relevant school staff.

### Participants

A total of 168 children were invited to participate in this study (n= 168; 87 male and 81 female), aged 8-9 years (mean age = 8.71, SD = 4.15). All participants were attending 3 different mainstream schools in the Bedfordshire county of the United Kingdom, and all had normal or corrected to normal vision and hearing. Children with a history of neurological or psychiatric conditions were excluded (i.e. severe neurodevelopmental delays, severe dyslexia and co-occurring autism and ADHD). Three schools agreed to take part in the study and implemented the intervention within their daily school program (Icknield Lower

School participated between September 5^th^ and December 6^th^, 2024; Parklea Primary School, and Leagrave Primary School participated between December 9^th^, 2024, and April 4^th^, 2025). Primary caregivers also completed a background questionnaire providing basic demographic information (child’s age, sex, any hearing or vision impairments). **Table 1** summarizes the demographic characteristics of the participant groups.

### Study Design

The study used a randomized controlled trial design. Enrollment was done at the class level, with all children given the opportunity to play (**Figure 1**) a custom digital intervention created by Mastermind Cognitive Training™ or our curated control battery of freely available math and coding applications (Kahoot!Multiplication, Sushi Monster, Dance Party, ScratchJr) ) that served as the control arm of this study. The Mastermind intervention was presented in VR via a Meta Quest 2™ or on a tablet via an Apple iPad™ (**Figure 2a-c**), while the control was presented only in tablet form (**Figure 2d**). Participants were randomly assigned to one of six groups: VR full dose (n = 28), VR half dose (n = 27), Tablet full dose (n = 28), Tablet half dose (n = 28), Control full dose (n = 23), Control half dose (n = 24). Training doses consisted of 25 minutes per training session for a full dose or 12.5 minutes for a half dose. The complete protocol is available in the Supplementary Materials (**Supplementary** Figure 5).

Students completed the training during the school day at a consistent time each week; make-up days were provided each week to allow those students who missed a training day to catch-up on their training sessions. Teachers at school were blinded as to which intervention and dosage were being given to their students. All participants were asked to participate in 30 training sessions over a period of 10 weeks (approximately 12.5 (full) or 6.25 (half) hours of training for each dosage assigned), and with the outcome measures conducted as pre- and post-training assessment sessions. Each student (regardless of group) was informed that their assigned training protocol was designed to train cognitive control abilities, using the same dialogue to avoid expectancy differences between groups. Students were encouraged to avoid discussing their training protocol with others to avoid potentially biasing participants in the other groups. After the 1^st^ day of training, students were asked to complete an expectancy survey^90^ to evaluate their impressions of how much improvement they expected on the objective outcome measures of attention (Right Eye, mCPT), reading and math fluency (SEA), and how enjoyable they believed their assigned intervention would be.

Participants originally assigned in the tablet group conditions were also offered the opportunity to complete a trial session of the Mastermind Cognitive Training program on the VR platform after the data collection was complete if they wished to do so. This was arranged as a ‘bonus and fun session’ in collaboration with the participating schools. All participants received a Certificate of Participation for their participation. The authors affirm that participants provided informed consent for publication of photographs taken during study activities for illustration purposes.

### Intervention

**Mastermind Cognitive Training**. The Mastermind Cognitive Training^TM^ program (**Figure 2)** integrates adaptive cognitive algorithms in a ‘gamified’ format that incorporates high-level 3D video game elements (e.g., art, music, story) to target cognitive and eye-tracking abilities. The training adjusts in real-time through a continuous feedback loop, providing feedback and rewards to create an engaging and interactive experience between the player and the game environment.

In each case, participants used either VR hand-held controllers (**Figure 2a**) or tapping on the tablet screen to respond to presented stimuli, in some cases ‘shooting’ specific target stimuli while avoiding others (**Figure 2b)**, or trying to catch colored targets in a given order while avoiding non-targets (**Figure 2c)**.

The cognitive training program consists of 5 modules, each focusing on a distinct aspect of cognitive control: a multiple object-tracking module, a visual search module, a working memory module, a rhythmic processing module, and eye exercises. Each module comprises adaptivity parameters where difficulty scales on a trial-by-trial basis, with a correct trial performed within a threshold-determined response window leading to increased difficulty on the following trial, and an incorrect trial leading to decreased difficulty on the following trial.

The **multiple object-tracking module** involves dynamic components aimed at engaging attention (via multiple object tracking), inhibitory control, and visual processing speed, with elements of increasing distraction. Participants are required to track multiple moving targets amongst distractors, over short block trials. The module uses evolving cued information combined with distractors, with the volume and speed of both cues and distractors increasing after correct trials and decreasing after incorrect trials.

The **visual search module** requires active scanning of the information presented on the screen in search of specific targets, resembling traditional visual search tasks. There are also task switching requirements, challenging cognitive flexibility resources that require participants to rapidly switch their focus based on distinct rules. The presented probes have features that integrate multiple rule bases, generating greater cognitive demands, similar to interference generated by a classic Stroop task^91,92^. Cued information is constantly evolving alongside numerous incongruent distracting features that increase as participant performance improves. For example, probes can appear in random locations across the screen, increasing the cognitive demands further by requiring rapid visual search and target identification, aiming to augment multitasking performance. This module also requires directional tracking (up, down, left, right), often aided by the brief presence of a directional cue indicating the screen location the target will appear while non-target elements are constantly appearing in an attempt to distract the user. This involves a constantly evolving amount of cued information as well as number of distracting elements, such that participants experience incrementally increasing cued and distractor elements as they advance and decreasing elements if they are incorrect in a trial.

The **working memory module** involves spatial working memory span engagement, comparable to the Corsi block task^93,94^. This requires participants to memorize object location when on screen, followed by a 5-7 second delay period after which participants are required to indicate where objects were located when previously visible. Correct responses lead to a greater number of targets displayed that participants would have to memorize on subsequent trials, while incorrect responses would reduce the number of targets until performance levels could be further challenged. Responses were provided using the controllers (VR) which feature as ‘punching gloves’ or tapping on the screen (Tablet). Participants are asked to memorize stimuli, with the addition of more stimuli or decreasing review and/or response windows after correct responses, or the subtraction of stimuli or increasing review and/or response windows after incorrect responses. Targets are presented as colors, shapes, number digits, and some iterations require hand-color coordinated response, thus increasing the spatial working memory load.

The **rhythm training module** (Rhythm IQ), derived from the software called Coherence at UCSF Neuroscape (see Zanto et al., 2024 for a detailed description), involves timed responses to stimuli presented in rhythmic sequences. This is a rhythm-based executive function task designed to enhance attention, working memory, and temporal processing^95,96^, promoting a synchronous tapping movement with the musical beat, replicating a ‘drumming-style’ performance. Participants use the controllers resembling ‘drumsticks’ (VR) or tap the screen (Tablet) in synchrony with the musical “beat”. This is also visually cued by moving orbs, with participants instructed to tap the target region once the orb fully overlaps it, corresponding with the musical beat. Rhythm and timing abilities are further challenged by the additional elements of ‘invisible rounds’ where the visual cues disappear and participants need to follow the beat pattern learned in previous rounds. Adaptive algorithms also modulate stimulus complexity and tempo to maintain optimal challenge levels for participants. Task difficulty is dynamically adjusted based on performance, increasing cognitive load by introducing faster rhythms, more complex stimulus-response mappings, and increased distractor interference. The rhythm training module adapts to correct/incorrect trial responses by adjusting the quantity and visibility of the cues, all while maintaining a rhythmic pattern that matches the beat of a song. Correct trial responses result in increased cues and patterns, both visible and invisible, while incorrect trial responses result in decreased cues and patterns.

The **eye exercises module** employs visual training techniques, including saccadic and smooth pursuit tracking and visual flexibility (near-far focusing). This is designed to enhance oculomotor control (eye movement, strengthening), visual attention, and eye–brain coordination^54,97–99^. This training is designed to reduce visual fatigue, improve fixation stability, and support cognitive functions such as spatial awareness and sustained attention^100^. This module consists of short sessions (30 seconds – 1.5 minutes) and is intended to complement the cognitive training protocol by optimizing the efficiency of visual input processing^101,102^. This module does not have adaptive trials based on accuracy of responses. While there are variations in trials across the same exercises, its random assignment of direction and the difficulty level does not change.

Each training session consisted of 3 eye exercises and 2 training protocol games. The rhythm training module was prescribed as the sole game played on every third training session.

Therefore, each training week followed this protocol: Session 1: eye exercises, game, game; Session 2: eye exercises, game, game; Session 3: eye exercises, game.

### Control Training

For the control arm of this study, we used four freely available math and coding applications (see **Figure 2**):

**Kahoot! Multiplication**

(https://kahoot.com^103,104^) is a game-based learning platform that enables users to engage in interactive and competitive gameplay. Players engage by tapping the color and shape on their device that corresponds to the answer they think is correct and earn points for responding quickly and accurately.

**Sushi Monster** is a gamified educational application developed by Scholastic and Blockdot to support learning of basic addition and multiplication facts^105,106^. Participants matched number-labelled sushi plates to target sums, encouraging flexible computation strategies and fact retrieval through game-based repetition^107^.

**Dance Party** is a visual coding activity developed by Code.org (https://code.org/dance) that uses block-based commands that animate dance routines, designed to build thinking skills and promote creativity and sequencing.

**ScratchJr** is a tool that enables users to create interactive stories and animations, while it supports storytelling, creative design, and logical reasoning. Participants created images and stories by dragging and dropping blocks on to the main work-frame on the device screen, promoting creative thinking and coding abilities^108,109^.

Implementation of these educational platforms mirrored the training process of the intervention groups. Participants in the Control groups were provided with tablets at the start of each session that had only these four applications enabled. All tablet devices used had internet access disabled so all math and coding applications used offline versions (as some of these applications, e.g. Kahoot! Multiplication, also support real-time interactive gameplay when internet-enabled). Participants used one of these applications for approximately 5-minute blocks and then switched to a different one, following this process for the duration of their session.

### Outcome Measures

**SEA:** Objective behavioral measures of academic performance were assessed via a custom application called SEA^20,59,60^ assessing basic academic performance (reading, math) at baseline (pre-training), and following the intervention period (post-training). This outcome measure is a computerized reading and math fluency test, comparable to the Woodcock-Johnson IV Tests of Achievement (WJ4; Sentence Reading Fluency and Math Facts Fluency subtests)^110^. The reading fluency assessment involved the presentation of short sentences (e.g. ‘A butterfly has ten wings’, ‘Ice is hot’, ‘An apple is blue’) and participants were instructed to respond by indicating whether the sentence presented was true or false using the computer keyboard. Participants were given 3 minutes to complete this assessment. The math fluency assessment involved simple arithmetic problems of addition, subtraction, and multiplication (e.g. 2 + 3, 6 – 1, 2 x 4) arranged in two tasks: a. participants would indicate their responses by selecting true or false (similar to the reading fluency task), and b. would use a number keyboard featured on the bottom of the screen to enter their responses. Participants were given 3 minutes to complete each of the math fluency tasks, thus, a total 9 minutes to administer. A rate correct score metric^20,111^ was used for each domain, taking into account the number of correct responses in 3 minutes while controlling for the response time taken for each correct measure.

**RightEye:** This outcome measure assessed eye-tracking behavior using the RightEye eye-tracking technology^112^. The RightEye assessment utilizes high-frequency infrared eye-tracking to objectively measure oculomotor behavior and visual processing. It is particularly suited for use with children in educational settings due to its short duration (∼5–7 minutes), child-friendly interface, and lack of need for verbal responses or advanced reading skills. It assesses various domains of eye movement and visual performance. The main output domains in the RightEye assessment that contribute to the RightEye overall score are: fixation stability (the ability to maintain steady gaze on a single point), saccadic eye movements (rapid eye movements between two points of focus), smooth pursuits (eye movements used to smoothly follow a moving object). In each case, a higher score is associated with better ocular performance.

**NICHQ Vanderbilt Assessment – Teacher Informant Scale:** The Vanderbilt Attention Deficit/Hyperactivity Disorder Rating Scale (VADPRS), which utilizes information based on the Diagnostic and Statistical Manual of Mental Disorders, 4th Ed. (DSM-IV), and was administered here to primarily assess changes in teacher perception of inattention as in our other work^9,23,24^. This measure was collected from the participants’ teacher prior to and immediately following the intervention period (approximately 10 weeks later). When completing this instrument, teachers were instructed to “please think about this student’s behaviors in the past 6 months, or since the last time this assessment was given.” Inattention concerns were assessed using the 1st 9 questions on the Vanderbilt, where participants’ teachers rated questions of inattention on a scale from 0-3, with 0 representing ‘never having a concern’, 1 ‘having occasional concerns’, 2 ‘often having concerns’, and 3 representing ‘very often having concerns’. The metric of interest for the Vanderbilt Assessment was calculated as being the total score across all questions presented as directed by the Vanderbilt Scoring Schema.

**Continuous Performance Task:** Our objective measure of attention, the mobile continuous performance task (mCPT) was derived from a well-validated custom continuous performance task, the Test of Variables of Attention (T.O.V.A.)^113^. Here participants maintain fixation on a central crosshairs and red squares are shown on a light grey background at the top or bottom of the field of view. During the infrequent condition, target stimuli were presented at the top of the screen as a 1:4 ratio of targets to nontargets and participants were instructed to only respond to these target stimuli. During the frequent condition, target stimuli were presented at the top of the screen as a 4:1 ratio of targets to nontargets and participants were instructed to only respond to these target stimuli. Participants completed 2 blocks of 125 trials for each condition. Our primary condition of interest was the frequent condition, with the variable of interest being response time variability (RTV), as this combination has been shown to be sensitive to changes following a digital intervention^8,10,21,114^.

**Mastermind Cognitive Assessments:** Each participant completed an assessment version of each Mastermind Cognitive training module (e.g. multiple object-tracking, visual search, and working memory). Participants completed one four-minute block in each case, and we specifically examined performance via the maximum level reached on each module. An overall MM assessment score was then calculated by averaging across all modules.

**Basic Response Time (BRT):** Finally, we also collected a control measure in the form of a basic response time (BRT) task to help evidence that any improvements on the cognitive measures were specific to attention and working memory processes and not simply the result of a general increase in basic speed of processing. In this task, participants responded to a target stimulus (40 trials) with a button press, first using their dominant hand and repeated using their less dominant hand. The task was to press the trigger on the VR controller or tap the Tablet screen (depending on group allocation) at the designated ‘button’ spot, as soon as the target red circle would appear on the screen. Here we assessed response time variability in line with our previous work^8,12^.

**Statistical Analysis:** All outcome measures were assessed with repeated measures ANCOVAs comparing pre-to post-training performance, with post-training performance serving as the dependent variable, pre-training performance serving as the covariate of interest, and group as the fixed factor of interest^115^. The use of ANCOVAs adjusts post-intervention scores based on individual baseline differences, which is critical when dealing with study designs such as this one. This is especially pertinent given that individuals who trained in VR were assessed in VR (and vice versa for those who trained on tablets), which could foster between-platform differences on the BRT, mCPT, and Mastermind tasks.

Planned follow-up t-tests and the Greenhouse-Geisser correction were used where appropriate. Spearman (nonparametric) correlations were used where appropriate. All statistical analyses were conducted using SPSS 24.0 (SPSS Inc.), with a p-value of .05 set as the threshold for significance. Critically, no between-group differences across all 6 groups were observed at baseline when assessed using one way ANOVA involving: (1) age (F_(5,151)_= 0.15 p= 0.98; (2) Vanderbilt inattention score (F_(5,153)_= 1.11 p= 0.36; (3) Right Eye Overall (F_(5,153)_= 0.71 p= 0.61; (4) SEA reading fluency (F_(5,154)_= 1.14 p= 0.34); (5) SEA math fluency (F_(5,147)_= 0.31 p= 0.91); (6) mCPT (F_(5,153)_= .674 p= 0.64); (7) Basic Response Time (F_(5,147)_= 1.74, p= 0.130).

## Data Availability Statement

The datasets generated during and/or analyzed during the current study are available from the corresponding author upon reasonable request.

## Code Availability Statement

The code used in the analysis are available from the corresponding author on reasonable request.

## Acknowledgements

We would like to thank all the participating teachers, school administrators, students, and parents from each school for being a part of this study, as well as the team of research coordinators (P Rzeszotarska, N Ahmed, T Tahat, I Giannikou, T Piasek, D Dimitrova) who helped with data collection for this study. Thanks to M Kloss and the team at Alternova for assistance with software testing and providing support throughout the study.

## Author Contributions

J.A.A, A.G, DF, MS designed the experiments; J.A.A., DF, MS developed the BBT software; J.A.A, A.G., DF, MS collected the data; J.A.A., A.G., C.G. and A.M.G. analyzed the data; and J.A.A., A.M.G, A.G. wrote the paper. All authors discussed the results.

