## Supplementary Materials for "Virtual Reality and Tablet Cognitive Training Improve Attention and Academic Skills Without Dose Effects"

[illegible]

| | | Mastermind Intervention | | Expectancy Matched Control | |
|---|---|---|---|---|---|
| | | Pre-Training Mean (SD) | Post-Training Mean (SD) | Pre-Training Mean (SD) | Post-Training Mean (SD) |
| Teacher Report | Vanderbilt Inattention Score | 6.15 (7.00) | 4.08 (5.13) | 6.72 (7.3) | 5.97 (7.0) |
| mCPT | RTV | 174 (100) | 129 (69) | 200 (107) | 171 (90) |
| SEA | Reading Fluency Efficiency Score | 0.31 (0.10) | 0.39 (0.12) | 0.30 (0.04) | 0.37 (0.08) |
| | Math Fluency Efficiency Score | 0.19 (0.06) | 0.21 (0.05) | 0.18 (0.10) | 0.20 (0.12) |
| MM | Overall Score | 9.77 (4.22) | 13.84 (2.54) | 9.73 (4.51) | 11.14 (4.28) |
| Right Eye | Overall Score | 70.3 (13.2) | 77.5 (10.5) | 69.5 (13.3) | 72.1 (11.6) |
| Basic Response Time | RTV | 186 (97) | 128 (92) | 182 (81) | 156 (93) |

[illegible]

| | | Mastermind **VR** Intervention | | Mastermind **Tablet** Intervention | |
|---|---|---|---|---|---|
| | | Pre-Training Mean (SD) | Post-Training Mean (SD) | Pre-Training Mean (SD) | Post-Training Mean (SD) |
| Teacher Report | Vanderbilt Inattention Score | 6.00 (7.29) | 3.96 (5.50) | 6.30 (6.77) | 4.20 (4.80) |
| mCPT | RTV | 175 (98) | 108 (54) | 173 (102) | 147 (76) |
| SEA | Reading Fluency Efficiency Score | 0.30 (0.11) | 0.38 (0.11) | 0.31 (0.10) | 0.39 (0.13) |
| | Math Fluency Efficiency Score | 0.18 (0.07) | 0.22 (0.05) | 0.19 (0.06) | 0.21 (0.06) |
| MM | Overall Score | 6.86 (3.40) | 12.90 (2.81) | 12.4 (2.91) | 14.8 (1.89) |
| Right Eye | Overall Score | 72.4 (11.86) | 79.7 (9.32) | 68.2 (14.4) | 75.5 (10.6) |
| Basic Response Time | RTV | 212 (100) | 154 (99) | 162 (88) | 106 (80) |

[illegible]

| | | VR FULL | | VR HALF | | Tablet FULL | | Tablet Half | |
|---|---|---|---|---|---|---|---|---|---|
| | | Pre-Training Mean (SD) | Post-Training Mean (SD) | Pre-Training Mean (SD) | Post-Training Mean (SD) | Pre-Training Mean (SD) | Post-Training Mean (SD) | Pre-Training Mean (SD) | Post-Training Mean (SD) |
| Teacher Report | Vanderbilt Inattention Score | 5.75 (7.4) | 4.92 (6.8) | 6.28 (7.3) | 2.88 (3.3) | 7.53 (7.5) | 5.21 (5.2) | 5.03 (5.8) | 3.14 (4.1) |
| mCPT | RTV | 157 (80) | 100 (40) | 193 (110) | 116 (64) | 169 (87) | 149 (75) | 177 (117) | 145 (78) |
| SEA | Reading Fluency Efficiency Score | .32 (.12) | .40 (.13) | .29 (.09) | .36 (.09) | .33 (.10) | .41 (.13) | .29 (.10) | .38 (.12) |
| | Math Fluency Efficiency Score | .18 (.08) | .22 (.05) | .19 (.06) | .21 (.05) | .19 (.06) | .20 (.05) | .19 (.06) | .21 (.06) |
| MM | Overall Score | 7.58 (4.1) | 13.94 (2.6) | 6.14 (2.3) | 11.67 (2.5) | 11.6 (3.3) | 14.9 (1.8) | 13.3 (2.2) | 14.7 (2.0) |
| Right Eye | Overall Score | 70.8 (13.4) | 80.7 (8.9) | 74.0 (9.9) | 78.8 (9.8) | 67.4 (15.2) | 74.9 (12.0) | 69.0 (13.6) | 76.0 (9.1) |
| Basic Response Time | RTV | 222 (100) | 137 (95) | 204 (100) | 170 (101) | 164 (95) | 100 (43) | 160 (83) | 111 (106) |

#### Supplementary Figure 1: Right Eye and Mastermind individual tasks (Hypothesis 1)

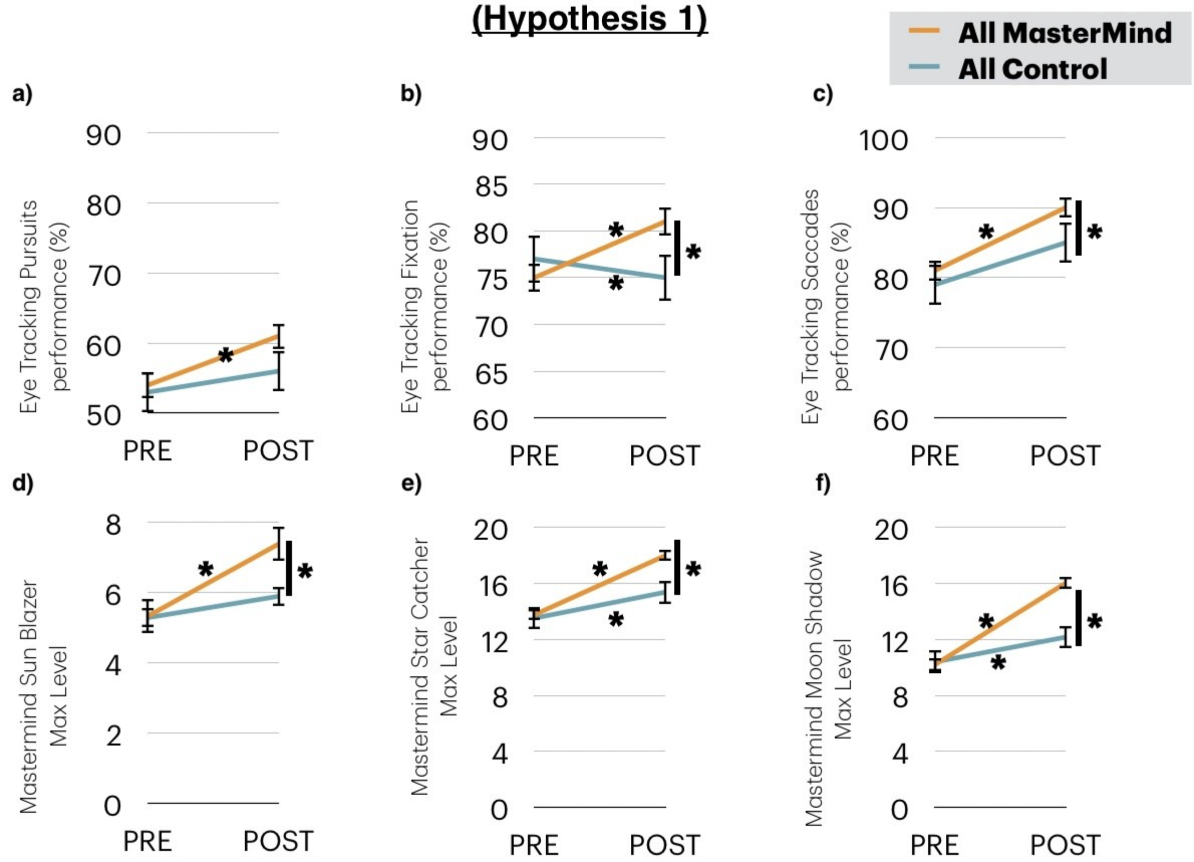

[illegible]. Panels of each group comparison for Hypothesis 1 for each measure. * = p< .05, |* = significant ANCOVA

#### Supplementary Figure 2: Right Eye and Mastermind Individual task for Hypothesis 2

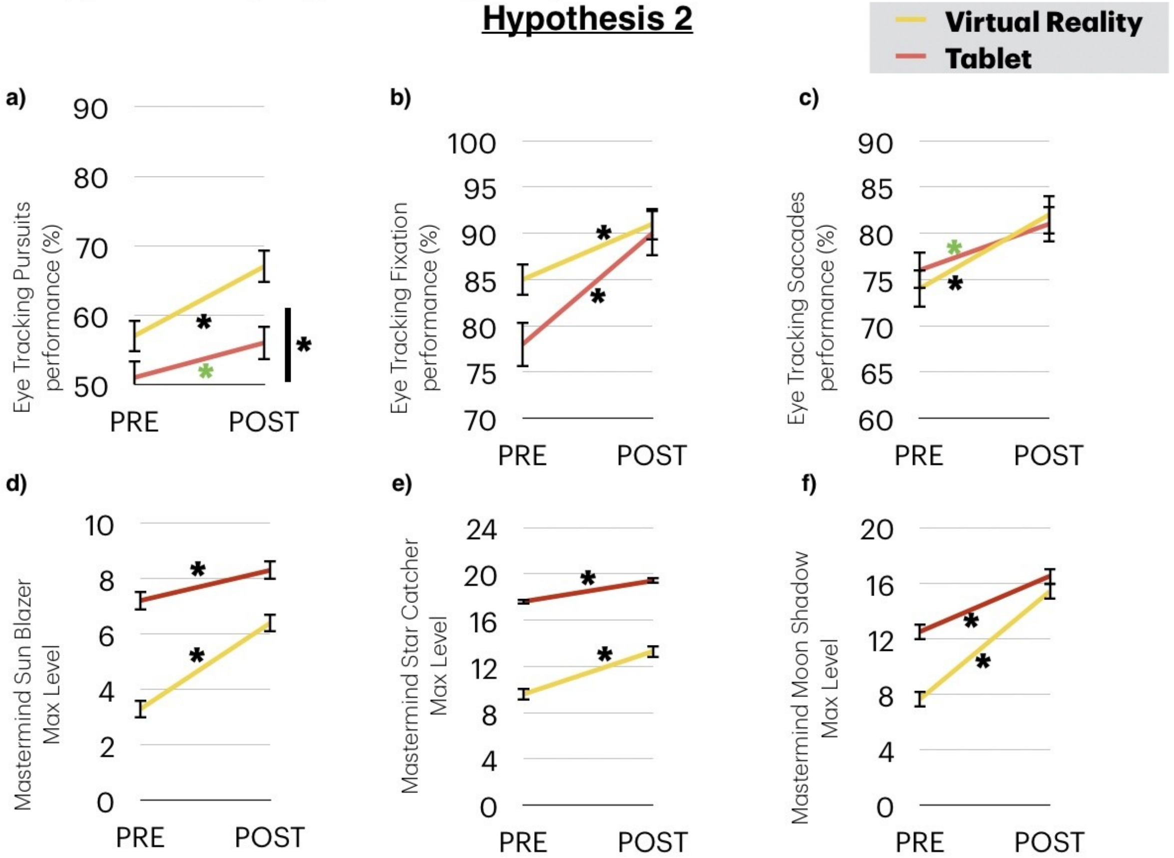

[illegible]. Panels of each group comparison for Hypothesis 1 for each measure. * = p< .05, |* = significant ANCOVA, Green * p < .07 (trend).

#### Supplementary Figure 3: Right Eye and Mastermind individual tasks for Hypothesis 3

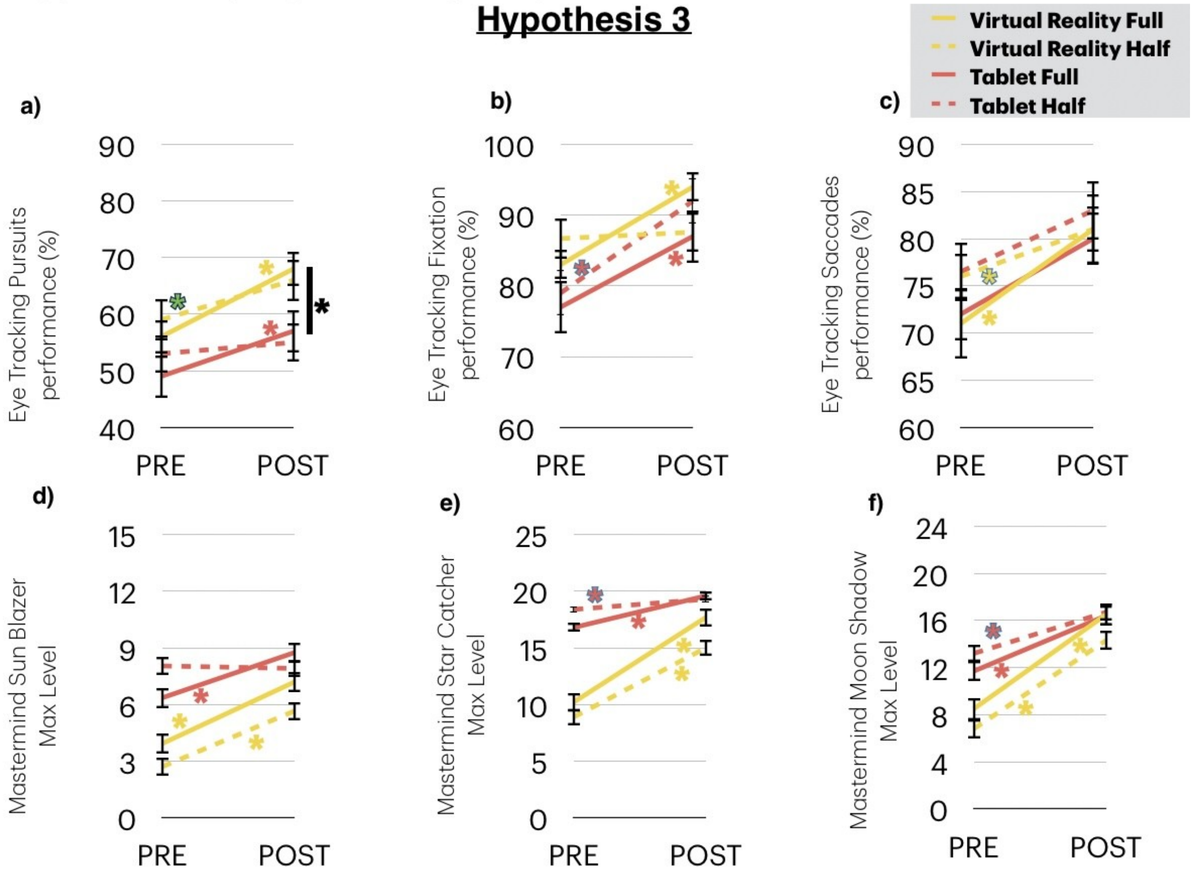

[illegible]. Panels of each group comparison for Hypothesis 1 for each measure. * = p< .05, |* = significant ANCOVA. Green * p < .07 (trend).

[illegible]: MM and Right Eye Individual Modules

| | | Mastermind Intervention | | Expectancy Matched Control | |
|---|---|---|---|---|---|
| | | Pre-Training Mean (SD) | Post-Training Mean (SD) | Pre-Training Mean (SD) | Post-Training Mean (SD) |
| MM Max Level | MM: Sunblazer | 5.32 (3.2) | 7.38 (2.5) | 5.28 (3.3) | 5.89 (3.2) |
| | MM: Star Catcher | 13.74 (5.5) | 17.96 (3.1) | 13.53 (6.1) | 15.38 (5.2) |
| | MM: Moon Shadow | 10.18 (5.5) | 16.01 (4.1) | 10.38 (5.5) | 12.15 (5.7) |
| RightEye | Fixation | 81.80 (20.6) | 90.34 (12.2) | 78.78 (20.1) | 84.95 (18.1) |
| | Saccades | 75.72 (18.3) | 81.64 (12.0) | 77.28 (20.1) | 75.43 (18.0) |
| | Pursuits | 54.49 (19.0) | 61.71 (18.0) | 53.47 (19.2) | 56.95 (20.1) |

[illegible]: MM and Right Eye Individual Modules

| | | Mastermind **VR** Intervention | | Mastermind **Tablet** Intervention | |
|---|---|---|---|---|---|
| | | Pre-Training Mean (SD) | Post-Training Mean (SD) | Pre-Training Mean (SD) | Post-Training Mean (SD) |
| MM Max Level | MM: Sun Blazer | 3.29 (2.4) | 6.39 (2.5) | 7.20 (2.8) | 8.29 (2.2) |
| | MM: Star Catcher | 9.58 (4.5) | 16.38 (3.7) | 17.61 (3.1) | 19.43 (1.2) |
| | MM: Moon Shadow | 7.65 (5.2) | 15.40 (4.2) | 12.52 (4.6) | 16.75 (3.9) |
| RightEye | Fixation | 85.63 (14.7) | 91.07 (11.8) | 77.98 (24.6) | 89.65 (12.5) |
| | Saccades | 74.92 (18.5) | 81.94 (13.1) | 76.52 (18.3) | 81.36 (11.1) |
| | Pursuits | 57.67 (18.4) | 67.46 (16.2) | 51.30 (19.3) | 56.29 (18.2) |

[illegible]: Outcome Measures for MM and RightEye

| | | VR FULL | | VR HALF | | Tablet FULL | | Tablet Half | |
|---|---|---|---|---|---|---|---|---|---|
| | | Pre-Training Mean (SD) | Post-Training Mean (SD) | Pre-Training Mean (SD) | Post-Training Mean (SD) | Pre-Training Mean (SD) | Post-Training Mean (SD) | Pre-Training Mean (SD) | Post-Training Mean (SD) |
| | MM: Sun Blazer | 3.92 (2.8) | 7.16 (2.4) | 2.69 (1.7) | 5.65 (2.3) | 6.39 (2.8) | 8.75 (2.0) | 8.04 (2.7) | 7.81 (2.4) |
| MM Max Level | MM: Star Catcher | 10.27 (5.2) | 17.73 (3.3) | 8.88 (3.6) | 15.04 (3.6) | 16.79 (3.8) | 19.61 (0.9) | 18.43 (1.9) | 19.25 (1.4) |
| | MM: Moon Shadow | 8.46 (6.0) | 16.50 (3.9) | 6.85 (4.3) | 14.31 (4.2) | 11.79 (5.2) | 16.43 (3.9) | 13.25 (3.9) | 16.71 (4.0) |
| | Fixation | 84.25 (15.6) | 94.45 (5.9) | 87.07 (13.9) | 87.89 (15.1) | 77.07 (23.7) | 87.25 (15.6) | 78.92 (26.0) | 92.15 (7.8) |
| RightEye | Saccades | 73.21 (15.0) | 80.24 (15.6) | 76.70 (13.3) | 83.53 (10.9) | 76.89 (19.1) | 81.14 (10.8) | 76.14 (17.7) | 81.59 (11.6) |
| | Pursuits | 56.14 (13.6) | 68.26 (18.1) | 59.25 (18.9) | 66.63 (18.6) | 49.46 (21.1) | 57.42 (19.1) | 53.22 (17.4) | 55.11 (17.6) |

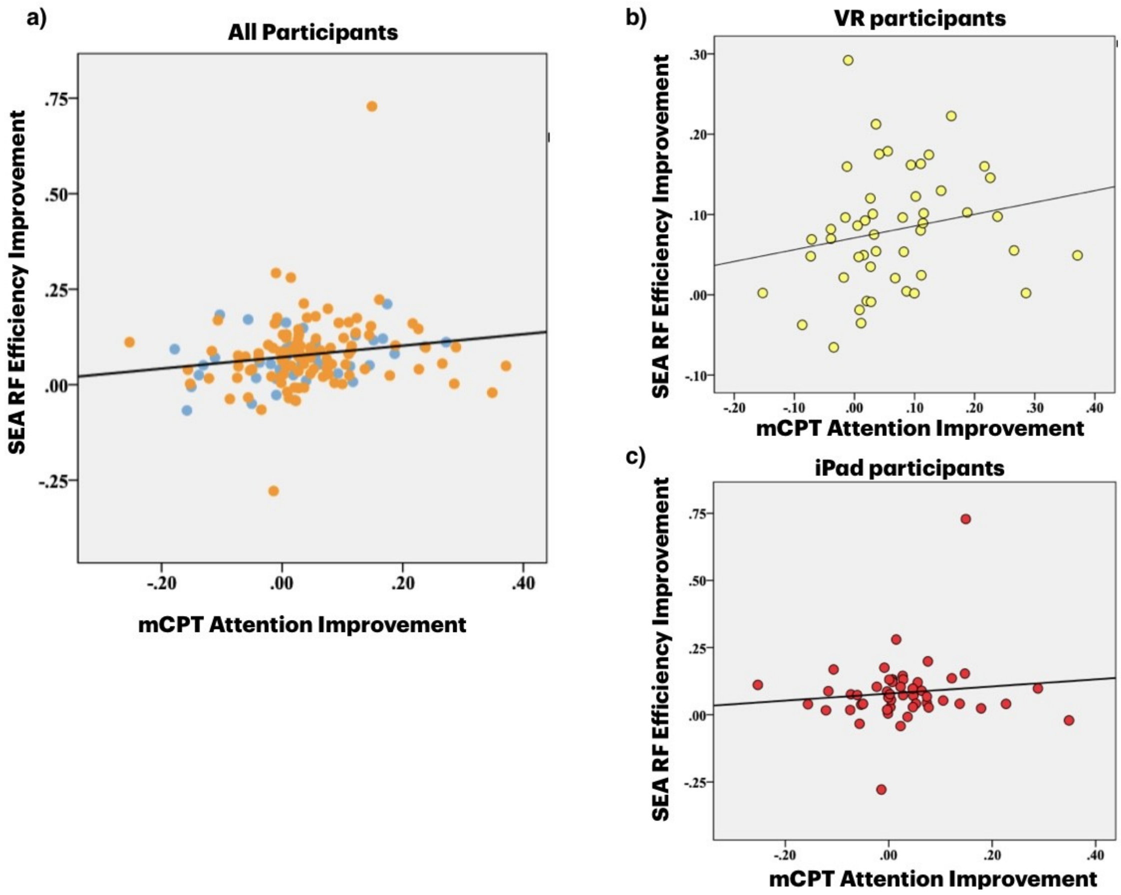

**[illegible].** Correlations between SEA Reading Fluency (RF) efficiency improvements and mCPT improvements on response time variability (RTV). a) Correlation across all participants ($r_{(143)}$= .22 p= .008). b) Correlation across all VR participants ($r_{(49)}$= .34 p= .018). c) Non-significant correlation across all iPad participants ($r_{(52)}$= .11 p= .44).

#### [illegible]

In the present study, we assessed academic performance using SEA, a custom application that offers a computerized reading and math fluency test, following the Woodcock-Johnson IV Tests of Achievement (WJ4; Schrank, Mather, & McGrew, 2014). We were also presented the opportunity to compare our SEA assessments of reading fluency to UK national standard teacher-administered assessments for each student. These measures of core academic skills help schools benchmark attainment and track student progress over time. More information about the UK National Curriculum for Key Stage 2 can be found at https://www.gov.uk/government/collections/national-curriculum.

Before and after the intervention phase, teachers assessed each student on reading performance based on UK national standard tests. These tests included metrics of reading comprehension and vocabulary. Teachers then coded student reading performance, first designating their grade of reading level (Year 1 – Year 4), and then whether they were working toward ('WTS'), at ('EXS'), or above ('GDS') age-related expectations within each age-based reading level. For example, 1WTS, 1EXS, and 1GDS, would refer to a student's reading level being equivalent to Year 1 and incremented based on the three designated descriptors (WTS, EXS, GDS), if reading level was equivalent to Year 2 would yield a descriptive score of 2WTS, 2EXS, or 2GDS. The same would apply for student reading level equivalent to Year 3 (3WTS, 3EXS, 3GDS) or Year 4 (4WTS, 4EXS, 4GDS). Student reading performance below this range would receive a descriptive score of ELG2, referring to early learning goals level 2. Therefore, the full scale used by the schools is: ELG2, 1WTS, 1EXS, 1GDS, 2WTS, 2EXS, 2GDS, 3WTS, 3EXS, 3GDS, 4WTS, 4EXS, 4GDS. We then transformed these codes to integers to reflect linear scale of performance for statistical analysis (ELG2 = 0, 1WTS = 1, 1EXS = 2, 1GDS = 3, 2WTS = 4, 2EXS = 5, 2GDS = 6, 3WTS = 7, 3EXS = 8, 3GDS = 9, 4WTS = 10, 4EXS = 11, 4GDS = 12). Given that the exact type of reading assessment differed between schools, we also conducted analyses with reading scores z-scored within school (in addition to the raw reading scores) to account for this potential difference. Here we observed a correlation between our SEA reading efficiency score and the UK reading scores collapsed across all participants both at pre-training ($r_{(141)} = .470$, $p = .0001$) and post-training ($r_{(139)} = .469$, $p = .0001$). When we tested each hypothesis using these scores in the same ANCOVA approach, we also observed the same pattern of results as with SEA: no group difference for Hypothesis 1 ($F_{(2,147)} = .65$ $p = 0.42$), Hypothesis 2 ($F_{(2,100)} = 0.058$ $p = 0.81$, and Hypothesis 3 ($F_{(4,98)} = 2.18$ $p = 0.14$).

#### [illegible]

The Mastermind Cognitive training would link to one of the 5 modules. The training protocol consisted of the following games:

1) Sun Blazer (SB) Game (with alternating game variations: Detect, Defend, Sequence, Symbol Change, Hear);

2) Star Catcher (SC) Game (with alternating game variations: Decide, Order, Switch);

3) Moon Shadow (MS) Game (with alternating game variations: Match, Reverse Order, Shape or Color);

4) RhythmIQ;

5) Eye exercises: Optokinetic, Saccades, Convergence/Divergence, Fixation, Near/Far.

#### [illegible]

Eye movement, control and strength are critical not only to vision, but also the ability to visually process information, focus, follow targets, and react. More specifically, the eye exercises included the following 5 tasks:

**Optokinetics:** This task involves compensatory reflex that supports visual stabilization. Participants had to watch as the ball steadily moved across the screen in various directions. They had to keep their head still throughout the task, follow the ball with their eyes and their task was to click the controller (VR) or tap the screen (Tablet) every time they saw the ball change colour. Participants were instructed to be sharp and quick as they only had a brief moment to react to the colour change.

**Saccades:** This task required rapid movement of the eyes between fixation points. An arrow within a ball that appeared at a variety of places around the screen, was pointing at a different direction on each trial. Participants were instructed to fixate on the arrow and use the joystick (VR) or swipe the screen (Tablet) to indicate the direction of the arrow.

**Convergence/Divergence:** This task is based on the ability to turn the eyes inward as an object moves closer (convergence) and outward as an object moves outward (divergence). Participants were instructed to track the ball using only their eyes while keeping their head still and quickly press the trigger button on their controllers (VR) or tap the screen (Tablet). They were instructed to be sharp and quick as they only had a brief moment to react to the color switch.

**Fixation:** This task required participants to maintain the gaze on a single object, specifically a ball that would change locations on the screen, while the background was in fluid motion. Participants were instructed to click the controller (VR) or tap the screen (Tablet) every time they saw the ball change color.

**Near / Far Shift:** This task is based on shifting focus from a near object(s) to a far object(s) quickly and continually. A ball with an arrow inside would appear at different points on the screen, either ‘near’ or ‘far’. Participants were instructed to keep tracking the arrow and use the joystick (VR) or swipe the screen (Tablet) to indicate the direction of the arrow.

#### [illegible]

Each game is structured to adapt dynamically to individual performance, with auto-leveling features that cater to the user's current skill level, scoring features and progression rewards, while challenging reaction time. Core skills trained were featured in the following tasks:

**Sun Blazer:** This task trained response inhibition processes. Participants had to shoot specific targets, often in a specific sequence, and avoid those that were 'traps'.

**Star Catcher:** This task trained multiple object tracking, with and without distraction elements, and instructions were presented to participants either auditorily only, or as targets featured on top of their screen. Hand-eye coordination was important as distractors had to be avoided and targets hit with precision in a fast-paced virtual environment.

**Moon Shadow:** This task trained working memory capacities through retention and recollection of information. Targets in the form of numbers, shapes, colors were presented in a gridded screen. Participants were allowed a few seconds to memorize the pattern, the targets would then disappear from the screen, specific targets would appear at the bottom of the screen and participants had to use their controllers (VR) to punch or tap the screen (Tablet) to indicate at which position the targets appeared. Different versions of this task featured targets as colors, shapes, numbers, either same or reverse series.

**RhythmIQ:** This task trained timing and rhythm abilities. As the music played, balls approached in synchronization with the song's beat pattern. Participants were instructed to use the controllers as 'drumsticks' (VR) or tap the screen (Tablet) to strike these balls with the corresponding hand's drumstick when they perfectly overlapped with the globes in front of the screen, following the song beat. Visibility Variation Rounds further challenged performance. For the Visible Rounds, the balls remained visible throughout, allowing participants to see and hit them in rhythm. For the Invisible Rounds, the balls would be invisible and participants had to continue to hit the globes based on the beat pattern they had learned but without the visual cues being available.

[illegible]

### VR TRAINING SESSIONS

| | **Eye Exercise 1** | **Eye Exercise 2** | **Eye Exercise 3** | **Training Game 1** | **Training Game 2** |
|---|---|---|---|---|---|
| Session 1 | Optokinetic | Saccades | Convergence/Divergence | SB - Sequence | SC - Order |
| Session 2 | Optokinetic | Fixation | Near/Far | MS - Match | SB - Hear |
| Session 3 | Optokinetic | Saccades | Convergence/Divergence | Rhythm IQ | |
| Session 4 | Optokinetic | Fixation | Near/Far | MS - Reverse Series | SC - Switch |
| Session 5 | Optokinetic | Saccades | Convergence/Divergence | SB - Detect | SC - Order |
| Session 6 | Optokinetic | Fixation | Near/Far | Rhythm IQ | |
| Session 7 | Optokinetic | Saccades | Convergence/Divergence | SB - Defend | MS - Number Order |
| Session 8 | Optokinetic | Fixation | Near/Far | SC - Order | MS - Shape or Color |
| Session 9 | Optokinetic | Saccades | Convergence/Divergence | Rhythm IQ | |
| Sesson 10 | Optokinetic | Fixation | Near/Far | SC - Listen & Switch | SB - Symbol Change |
| Session 11 | Optokinetic | Saccades | Convergence/Divergence | SB - Defend | MS - Shape or Color |
| Session 12 | Optokinetic | Fixation | Near/Far | Rhythm IQ | |
| Session 13 | Optokinetic | Saccades | Convergence/Divergence | SC - Decide | MS - Shapes |
| Session 14 | Optokinetic | Fixation | Near/Far | SB - Sequence | MS - Match |
| Session 15 | Optokinetic | Saccades | Convergence/Divergence | Rhythm IQ | |
| Session 16 | Optokinetic | Fixation | Near/Far | MS - Reverse Series | SC - Order |
| Session 17 | Optokinetic | Saccades | Convergence/Divergence | SB - Symbol Change | SC - Order |
| Session 18 | Optokinetic | Fixation | Near/Far | Rhythm IQ | |
| Session 19 | Optokinetic | Saccades | Convergence/Divergence | SC - Listen & Switch | SB - Detect |
| Session 20 | Optokinetic | Fixation | Near/Far | SB - Hear | MS - Number Order |
| Session 21 | Optokinetic | Saccades | Convergence/Divergence | Rhythm IQ | |
| Session 22 | Optokinetic | Fixation | Near/Far | SC - Decide | MS - Shape or Color |
| Session 23 | Optokinetic | Saccades | Convergence/Divergence | SB - Defend | MS - Match |
| Session 24 | Optokinetic | Fixation | Near/Far | Rhythm IQ | |
| Session 25 | Optokinetic | Saccades | Convergence/Divergence | SB - Defend | SC - Order |
| Session 26 | Optokinetic | Fixation | Near/Far | MS - Match | SB - Hear |
| Session 27 | Optokinetic | Saccades | Convergence/Divergence | Rhythm IQ | |
| Session 28 | Optokinetic | Fixation | Near/Far | MS - Shape or Color | SC - Switch |
| Session 29 | Optokinetic | Saccades | Convergence/Divergence | SB - Sequence | SC - Order |
| Session 30 | Optokinetic | Fixation | Near/Far | Rhythm IQ | |

**Key:**

SB = Sun Blazer Game (with alternating game variations: Detect, Defend, Sequence, Symbol Change, Hear)

SC = Star Catcher Game (with alternating game variations: Decide, Order, Switch)

MS = Moon Shadow Game (with alternating game variations: Match, Reverse Order, Shape or Color)

### TABLET TRAINING SESSIONS

| | Eye Exercise 1 | Eye Exercise 2 | Eye Exercise 3 | Training Game 1 | Training Game 2 |
|---|---|---|---|---|---|
| Session 1 | Optokinetic | Saccades | Convergence/Divergence | SC - Decide | SB - Detect |
| Session 2 | Optokinetic | Fixation | Near/Far | MS - Number Order | SC - Listen |
| Session 3 | Optokinetic | Saccades | Convergence/Divergence | Rhythm IQ | |
| Session 4 | Optokinetic | Fixation | Near/Far | SB - Time | MS - Reverse Series |
| Session 5 | Optokinetic | Saccades | Convergence/Divergence | SC - Order | SB - Hear |
| Session 6 | Optokinetic | Fixation | Near/Far | Rhythm IQ | |
| Session 7 | Optokinetic | Saccades | Convergence/Divergence | MS - Shape or Color | SC - Decide |
| Session 8 | Optokinetic | Fixation | Near/Far | SB - Symbol Change | MS - Number Order |
| Session 9 | Optokinetic | Saccades | Convergence/Divergence | Rhythm IQ | |
| Sesson 10 | Optokinetic | Fixation | Near/Far | SC - Switch & Decide | SB - Detect |
| Session 11 | Optokinetic | Saccades | Convergence/Divergence | MS - Shapes | SC - Track |
| Session 12 | Optokinetic | Fixation | Near/Far | Rhythm IQ | |
| Session 13 | Optokinetic | Saccades | Convergence/Divergence | SB - Hear | MS - Reverse Series |
| Session 14 | Optokinetic | Fixation | Near/Far | SC - Decide | SB - Time |
| Session 15 | Optokinetic | Saccades | Convergence/Divergence | Rhythm IQ | |
| Session 16 | Optokinetic | Fixation | Near/Far | MS - Shape or Color | SC - Listen |
| Session 17 | Optokinetic | Saccades | Convergence/Divergence | SB - Hear | MS - Shapes |
| Session 18 | Optokinetic | Fixation | Near/Far | Rhythm IQ | |
| Session 19 | Optokinetic | Saccades | Convergence/Divergence | SC - Order | SB - Symbol Change |
| Session 20 | Optokinetic | Fixation | Near/Far | MS - Number Order | SC - Decide |
| Session 21 | Optokinetic | Saccades | Convergence/Divergence | Rhythm IQ | |
| Session 22 | Optokinetic | Fixation | Near/Far | SB - Detect | MS - Reverse Series |
| Session 23 | Optokinetic | Saccades | Convergence/Divergence | SC - Switch & Decide | SB - Time |
| Session 24 | Optokinetic | Fixation | Near/Far | Rhythm IQ | |
| Session 25 | Optokinetic | Saccades | Convergence/Divergence | MS - Shape or Color | SC - Track |
| Session 26 | Optokinetic | Fixation | Near/Far | SB - Hear | MS - Shapes |
| Session 27 | Optokinetic | Saccades | Convergence/Divergence | Rhythm IQ | |
| Session 28 | Optokinetic | Fixation | Near/Far | SC - Decide | SB - Detect |
| Session 29 | Optokinetic | Saccades | Convergence/Divergence | MS - Number Order | SC - Listen |
| Session 30 | Optokinetic | Fixation | Near/Far | Rhythm IQ | |

### VR Instructions (assessments)

| NAME OF ASSESSMENT: | GOAL OF THIS ASSESSMENT PORTION: | INSTRUCTIONS: |
|---|---|---|
| Basic Reaction Time Assessment | **Response Time:** Discover how quickly you can react to stimulus. | 1. **Center Focus:** Keep your eyes locked on the cross at the screen's center throughout the round.<br>2. **Red Circle Alert:** Be on the lookout for a red circle that will appear at the screen's center periodically.<br>3. **Quick Trigger:** As soon as you see the circle, swiftly press the trigger on the controller with your dominant hand, using your pointer finger. The quicker the response, the better.<br>4. **Switch Hands:** The next round, switch to your non-dominant hand and repeat the same exercise. |
| Continuous Performance Assessment | **Focus And Attention:** Measure your ability to stay focused and attentive. | 1. **Steadfast Focus:** Throughout the round, keep your gaze fixed on the cross at the center of the screen<br>2. **Spot the Red Square:** Watch for a red square that will appear either at the top or bottom of the screen, near the central cross.<br>3. **Swift Response:** When you see the square at the top, react rapidly by pressing the trigger on either controller with your pointer finger. If it's at the bottom, hold off with no action.<br>4. **Repeat with Adaptability:** Do the same in a second round, using either hand, but only respond to the top red square. |
| Visual Signaling Assessment | **Visual Reaction:** Test your skill in reacting to visual cues while filtering out distractions. | 1. **Central Focus:** Throughout the round, keep your eyes locked on the cross at the center of the screen.<br>2. **Peripheral Vision Test:** A red triangle will pop up either to the left or right of the central cross. Use your peripheral vision to spot it, while maintaining focus on the cross.<br>3. **Quick Reflexes:** Match the triangle's side with your controller. If it appears on the right, press the right controller's trigger, and vice versa.<br>4. **Ignore Distractions:** Be wary of arrows that may mislead you. They're there to test your ability to stay focused under distracting conditions. |
| Multiple Object Tracking Assessment | **Visual Tracking:** Establish a baseline for how well you can track and respond to moving objects. | 1. **Gear Up:** Ready each hand with a paddle.<br>2. **Orb Challenge:** Imagine floating orbs as dynamic targets moving in all directions, testing your reaction time.<br>3. **Precision Strikes:** Use the trigger on your controller to hit targets with precision. Hold until you make contact.<br>4. **Use Both Hands:** Strike with either hand. It's about adaptability and quick thinking. |
| Identification and Inhibition Assessment | **Task-Switching:** Measure your agility in shifting between various tasks. | 1. **Arm Up with Blasters:** Equip a blaster in each hand using your Oculus controllers.<br>2. **Shape & Hand Coordination:** Look for shapes displayed at the top of the screen, each assigned to a specific hand.<br>3. **Target Precision:** Random targets, each with a shape, will appear on the screen. Strike only the targets with your assigned shape using the correct hand.<br>4. **Adaptive Challenge:** Stay alert as the shape and hand assignments change periodically. This tests your ability to adapt swiftly. |

| | | |
|---|---|---|
| Working Memory Assessment | **Visual Memory:** Gauge your memory skills and ability to utilize visual cues. | 1. **Gloved and Ready:** Put on a glove on each hand, using your Oculus controllers.<br>2. **Visual Formation Challenge:** A formation of boxes will appear, some black and others with yellow shapes.<br>3. **Memory Drill:** Remember which boxes contain which shapes.<br>4. **Pattern Pause:** The pattern will vanish briefly, testing your ability to retain information.<br>5. **Shape Assignment:** You'll then get a cue to hit a specific shape.<br>6. **Strategic Strikes:** Targets in the same box formation will approach you. Like a strategic move in a game, punch only the targets that match your shape assignment.<br>7. **Precision Punching:** Your task is to accurately punch the designated targets. |

### iPad Instructions (assessments)

| **NAME OF ASSESSMENT :** | **GOAL OF THIS ASSESSMENT PORTION:** | **INSTRUCTIONS:** |
|---|---|---|
| Basic Reaction Time Assessment | **Response Time:** Discover how quickly you can react to stimulus. | 1. **Center Focus:** Keep your eyes locked on the cross at the screen's center throughout the round.<br>2. **Red Circle Alert:** Be on the lookout for a red circle that will appear at the screen's center periodically.<br>3. **Quick Trigger:** As soon as you see the circle, swiftly tap the button in the bottom corner of your screen on your dominant hand side. The quicker the response, the better.<br>4. **Switch Hands:** The next round, switch to your non-dominant hand and repeat the same exercise. |
| Continuous Performance Assessment | **Focus And Attention:** Measure your ability to stay focused and attentive. | 1. **Steadfast Focus:** Throughout the round, keep your gaze fixed on the cross at the center of the screen<br>2. **Spot the Red Square:** Watch for a red square that will appear either at the top or bottom of the screen, near the central cross.<br>3. **Swift Response:** When you see the square at the top, react rapidly by tapping the button in the bottom corner of the screen. If the square is at the bottom, hold off with no action.<br>4. **Repeat with Adaptability:** Do the same in a second round, again only responding to the top red square. |
| Visual Signaling Assessment | **Visual Reaction:** Test your skill in reacting to visual cues while filtering out distractions. | 1. **Central Focus:** Throughout the round, keep your eyes locked on the cross at the center of the screen.<br>2. **Peripheral Vision Test:** A red triangle will pop up either to the left or right of the central cross. Use your peripheral vision to spot it, while maintaining focus on the cross.<br>3. **Quick Reflexes:** Match the triangle's side with the buttons in the bottom corner of your screen. If it appears on the right, press the right button, and vice versa.<br>4. **Ignore Distractions:** Be wary of arrows that may mislead you. They're there to test your ability to stay focused under distracting conditions. |
| Multiple Object | **Visual Tracking:** Establish a baseline for how well you | 1. **Orb Challenge:** Imagine floating orbs as dynamic targets moving in all directions, testing your reaction time by shooting in your direction. |

| | | |
|---|---|---|
| Tracking Assessment | can track and respond to moving objects. | 2. **Precision Strikes:** Tap all targets with precision. Strike with either hand. It's about adaptability and quick thinking |
| Identification and Inhibition Assessment | **Task-Switching:** Measure your agility in shifting between various tasks. | 1. **Shape & Hand Coordination:** Look for target assignment displayed at the top of the screen, Targets will be a specific shape and color.<br>2. **Target Precision:** Random targets will appear on the screen. Tap only the targets with your assigned shape and color.<br>3. **Adaptive Challenge:** Stay alert as the target assignments will change periodically. This tests your ability to adapt swiftly. |
| Working Memory Assessment | **Visual Memory:** Gauge your memory skills and ability to utilize visual cues. | 1. **Visual Formation Challenge:** A formation of boxes will appear, some black and others with yellow shapes.<br>2. **Memory Drill:** Remember which boxes contain which shapes.<br>3. **Pattern Pause:** The pattern will vanish briefly, testing your ability to retain information.<br>4. **Shape Assignment:** You'll then get a cue to tap a specific shape.<br>5. **Strategic Strikes:** Targets in the same box formation will approach you. Like a strategic move in a game, tap only the targets that match your shape assignment. |
